# A fetal oncogene NUAK2 is an emerging therapeutic target in glioblastoma

**DOI:** 10.1101/2024.12.31.630965

**Authors:** Hanhee Jo, Aneesh Dalvi, Wenqi Yang, Elizabeth Morozova, Sarah Munoz, Stacey M. Glasgow

## Abstract

Glioblastoma Multiforme (GBM) is the most prevalent and highly malignant form of adult brain cancer characterized by poor overall survival rates. Effective therapeutic modalities remain limited, necessitating the search for novel treatments. Neurodevelopmental pathways have been implicated in glioma formation, with key neurodevelopmental regulators being re-expressed or co-opted during glioma tumorigenesis. Here we identified a serine/threonine kinase, NUAK family kinase 2 (NUAK2), as a fetal oncogene in mouse and human brains. We found robust expression of NUAK2 in the embryonic brain that decreases throughout postnatal stages and then is re-expressed in malignant gliomas. However, the role of NUAK2 in GBM tumorigenesis remains unclear. We demonstrate that CRIPSR-Cas9 mediated NUAK2 deletion in GBM cells results in suppression of proliferation, while overexpression leads to enhanced cell growth in both *in vitro* and *in vivo* models. Further investigation of the downstream biological processes dysregulated in the absence of NUAK2 reveals that NUAK2 modulates extracellular matrix (ECM) components to facilitate migratory behavior. Lastly, we determined that pharmaceutical inhibition of NUAK2 is sufficient to impede the proliferation and migration of malignant glioma cells. Our results suggest that NUAK2 is an actionable therapeutic target for GBM treatment.

## INTRODUCTION

Glioblastoma (GBM) is the most common and lethal brain tumor (Aldape et al., 2019; Deorah et al., 2006). These devastating tumors exhibit widespread invasion throughout the brain, are highly proliferative, and are resistant to chemotherapy and radiotherapy (Stupp et al., 2005, 2009; Van Meir et al., 2010; Xu et al., 2020); making them exceedingly difficult to treat (Konishi et al., 2012; Louis et al., 2021; McDonald et al., 2011; Milano et al., 2010; Omuro & DeAngelis, 2013; Weller et al., 2015). Even with the current standard of care, including surgical resection, radiation, and chemotherapy, the prognosis for glioblastoma is dismal, with a median survival rate of 15 months (Verdugo et al., 2022; Weller et al., 2015). Therefore, identifying new efficient molecular targets is crucial for the development of therapeutic strategies.

Neurodevelopmental signaling pathways and transcriptional cascades have also been implicated in glioma tumor initiation, maintenance, and progression (Baker et al., 2016; Curry & Glasgow, 2021; Mehta, 2018; Sojka & Sloan, 2024). The growing literature defining roles for these developmental genes in tumorigenesis has revealed a subclass of oncogenes called fetal oncogenes, which are predominantly expressed during embryonic development and cancer, but their expression is nominal in adult tissues (Cao et al., 2023; West et al., 2018).

The minimal expression of fetal oncogenes in normal tissue can be exploited to allow for more precise targeting of cancer cells with marginal off-target effects of normal cells or neurotoxicity.

Tumor progression is controlled by molecular mechanisms triggered by multiple signaling pathways, often through the activation of regulatory kinases (Manning et al., 2002; Nakada et al., 2020; Schlessinger, 2000). Kinase activity directly affects the activation/inactivation of downstream effectors which are crucial for the initiation of many biological phenomena, such as cell growth, proliferation, and apoptosis (Fleuren et al., 2016). Mutations and alterations in several kinase signaling cascades have been associated with glioma tumorigenesis leading to inhibition of apoptosis, cellular proliferation, and tissue invasion (Aiello & Stanger, 2016; Balachandran & Narendran, 2023; Ma et al., 2010; Monk & Holding, 2001). The abnormal expression or activity of kinases, which can be specific to cancerous cells, represents an attractive target for glioma therapy (Adjei, 2005; Stitzlein et al., 2024).

NUAK family kinase 2 (NUAK2), also known as sucrose non-fermenting (SNF-1)-like kinase (SNARK), is a serine/threonine kinase of the AMP-activated protein kinase family. NUAK2 is crucial for the formation of the central nervous system (CNS) and has been shown to have a role in various solid tumors. In developing mice, NUAK2 expression is found in the neural folds, and NUAK2 knockout mice show neural tube closure defects, including exencephaly (Hirano et al., 2006; Ohmura et al., 2012). Similarly, loss-of-function mutations of NUAK2 in humans result in anencephaly, a severe form of neural tube closure failure (Bonnard et al., 2020). In both mice and humans, these neural tube defects are linked to defective regulation of cytoskeletal proteins (Bonnard et al., 2020; Ohmura et al., 2012). Roles for NUAK2 in several non-CNS cancers have been reported, with its expression being highly correlated with tumor progression and poor prognosis in patients (Chen et al., 2022; Fu et al., 2022; Li et al., 2021; Namiki et al., 2011, 2015; Tang et al., 2017; Wang et al., 2024). However, there is limited knowledge of the role of NUAK2 in GBM.

In this study, we find that NUAK2 is a fetal oncogene whose expression is low in juvenile and adult brains but high in developing brain and glioblastoma patients. In GBM cells, we show that NUAK2 deletion leads to attenuation of proliferation and migration, while overexpression enhances these processes. Modulation of NUAK2 expression in *in vivo* models of malignant glioma mimics these results. Importantly, pharmaceutical inhibition of NUAK2 exhibits significant effects in mitigating glioma progression. Therefore, NUAK2 is potential actionable target for the treatment of GBM.

## RESULTS

### NUAK2, a fetal oncogene, is associated with poor prognosis in GBM patients

NUAK2 plays a crucial role in brain development and the formation of non-CNS solid tumors, with its expression or mutations leading to various abnormalities. To investigate whether NUAK2 functions as a fetal oncogene in the CNS, we first examined publicly accessible RNA- sequencing (RNA-seq) data from the BrainSpan Atlas for the developing human brain, which profiles up to sixteen cortical and subcortical structures throughout the entire span of human brain development. Our analysis revealed that NUAK2 mRNA expression is significantly elevated during early developmental stages, declining gradually and remaining silent after birth in human brains (Fig 1A). In contrast, RNA-sequencing data from The Cancer Genome Atlas (TCGA) indicate that NUAK2 expression is markedly elevated in GBM patients, while normal brain tissues display only minimal expression levels (Fig 1B). Additionally, our TCGA analysis found a correlation between NUAK2 levels and glioma tumor grade, where high-grade gliomas exhibited greater NUAK2 expression than low-grade gliomas (Fig 1C and D); implying that NUAK2 may play a role in brain tumor malignancy.

**Figure 1.**
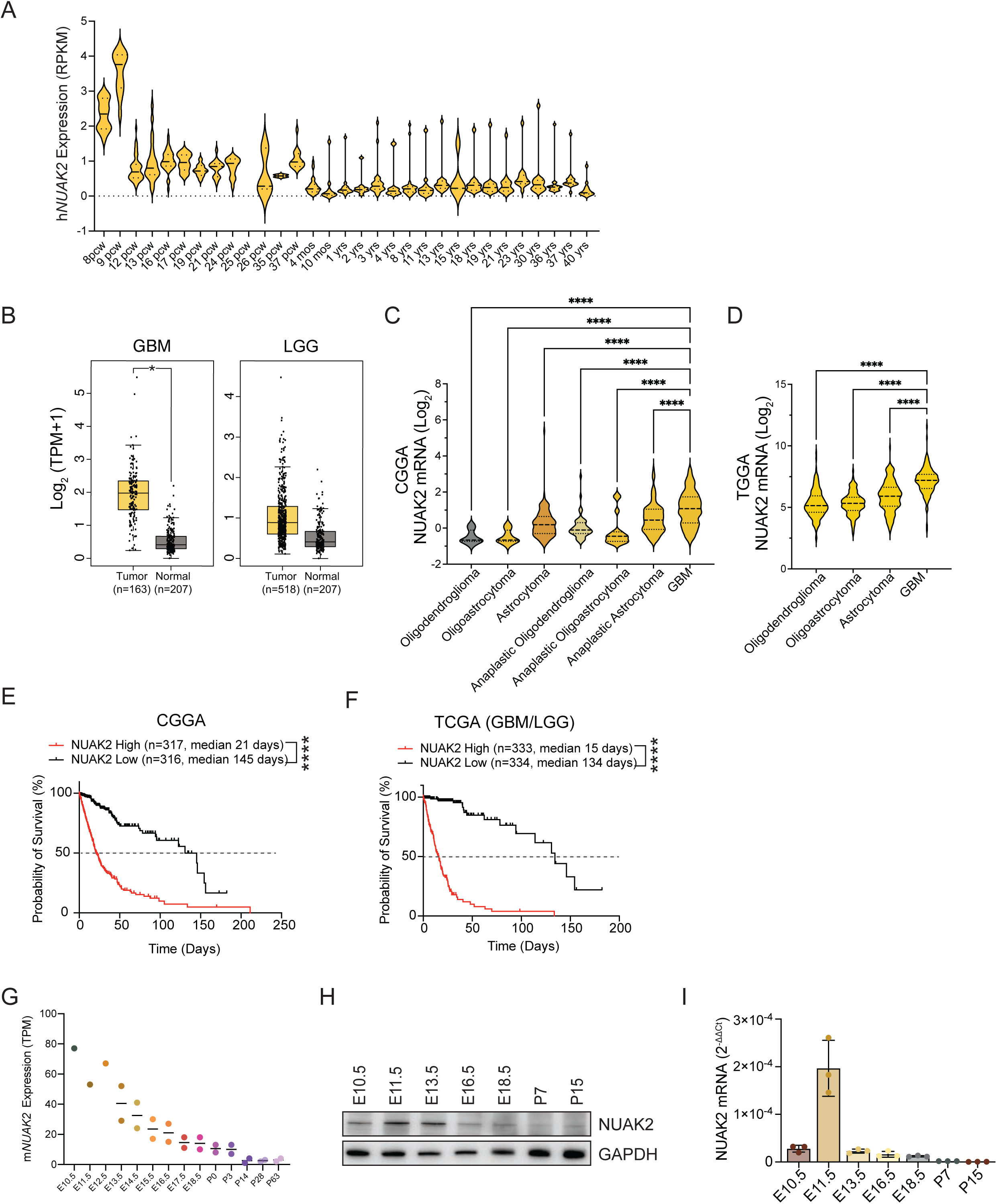
A fetal oncogene NUAK2 is associated with poor prognosis in GBM patients. **A** RPKM-normalized NUAK2 mRNA expression of various human brain regions from 8 post- conception weeks (pcw) to 40 years of age. Data was obtained from the BrainSpan Atlas. **B** Normalized NUAK2 mRNA expression of TCGA GBM (n = 163) or low-grade glioma (LGG) (n = 518) and GTEx non-tumor (n = 207) samples (*p < 0.05; Statistical significance was determined by One-way ANOVA). Shown are mean ±SD. Data were obtained from GEPIA (http://gepia.cancer-pku.cn/). **C** NUAK2 mRNA expression across glioma subtypes showing the highest expression in GBM in the CGGA dataset (****p < 0.0001; Statistical significance was determined by one-way ANOVA followed by Tukey’s multiple comparisons test). Data are represented as mean ±SD. **D** NUAK2 mRNA expression across glioma subtypes showing the highest expression in GBM in the TCGA dataset (****p < 0.0001; Statistical significance was determined by one-way ANOVA followed by Tukey’s multiple comparisons test). Data are represented as mean ±SD. **E** Kaplan-Meier survival analysis from CGGA of high (21 days; n = 317) and low (145 days; n = 316) NUAK2 expressors shows high NUAK2 expression is correlated with worse survival outcomes (***p < 0.001; Statistical significance was determined by log-rank (Mantel-Cox) test). **F** Kaplan-Meier survival analysis from TCGA of high (15 days; n = 333) and low (134 days; n = 334) NUAK2 expressors shows high NUAK2 expression is correlated with worse survival outcomes across low- and high-grade malignant gliomas (***p < 0.001; Statistical significance was determined by log-rank (Mantel-Cox) test). **G** TPM-normalized NUAK2 mRNA expression of mouse forebrain or hindbrain ranging from embryonic day 10.5 to postnatal day 63. Data was obtained from EMBL’s European Bioinformatics Institute (EMBL-EBI; https://www.ebi.ac.uk/). **H** Representative western blot of NUAK2 protein expression in whole wildtype (WT) embryonic brain tissue across 7 stages of development. GAPDH was used as a loading control. **I** Representative qRT-PCR of NUAK2 mRNA expression in WT embryonic brain tissues across developmental stages. Data was normalized to GAPDH (n=3).

To assess the relationship between NUAK2 expression and overall survival in human gliomas, we analyzed TCGA and The Chinese Glioma Genome Atlas (CGGA) datasets. Analysis of overall patient 50% survival rates revealed that elevated NUAK2 levels are strongly associated with reduced survival rates (Fig 1E and F). Together with our gene expression analysis, these findings suggest that NUAK2 functions as a fetal oncogene and demonstrates an explicit relationship between tumor progression and NUAK2 expression in GBM.

To further confirm our analysis, we examined NUAK2 expression in mice across different ages using publicly available datasets from EMBL’s European Bioinformatics Institution of developing mouse brain transcriptomes (Cardoso-Moreira et al., 2019). Similar to humans, NUAK2 mRNA expression in mice peaks during early development and declines postnatally (Fig 1G). Our immunoblot and qPCR analyses further confirmed this trend, with high NUAK2 levels in developing mouse brains, which substantially reduced in expression after birth (Fig 1H and I). Together, these findings classify NUAK2 as a fetal oncogene and demonstrate its strong association with GBM prognosis and tumor progression.

### NUAK2 is critical for GBM cell proliferation

To understand the role of NUAK2 in GBM, we investigated the impacts of loss-of-function (LOF) and gain-of-function (GOF) studies in GBM cells. mRNA and protein expression analysis across four GBM cell lines (U87, LN229, U251, and LN319) revealed varying NUAK2 levels, with U251 and LN319 showing high expression, LN229 moderate levels, while U87 cells exhibiting nominal NUAK2 expression (Fig 2A and B). To determine whether NUAK2 is essential for promoting glioblastoma cell growth, we silenced NUAK2 in U251 cells using a CRISPR-Cas9 system, as these cells have high NUAK2 expression (Fig 2A and B). Successful NUAK2 deletions were confirmed by both immunoblot and qPCR in three independent clones (Fig. 2C and D). Anti-Ki67 staining revealed significantly reduced cell proliferation in NUAK2- CRISPR (CR) cells (Fig 2E). Additionally, our proliferation (MTT) and colony formation assays showed that NUAK2 deletion resulted in reduced growth of glioblastoma cells and suppressed formation of colonies in the absence of NUAK2 (Fig 2F and G).

**Figure 2.**
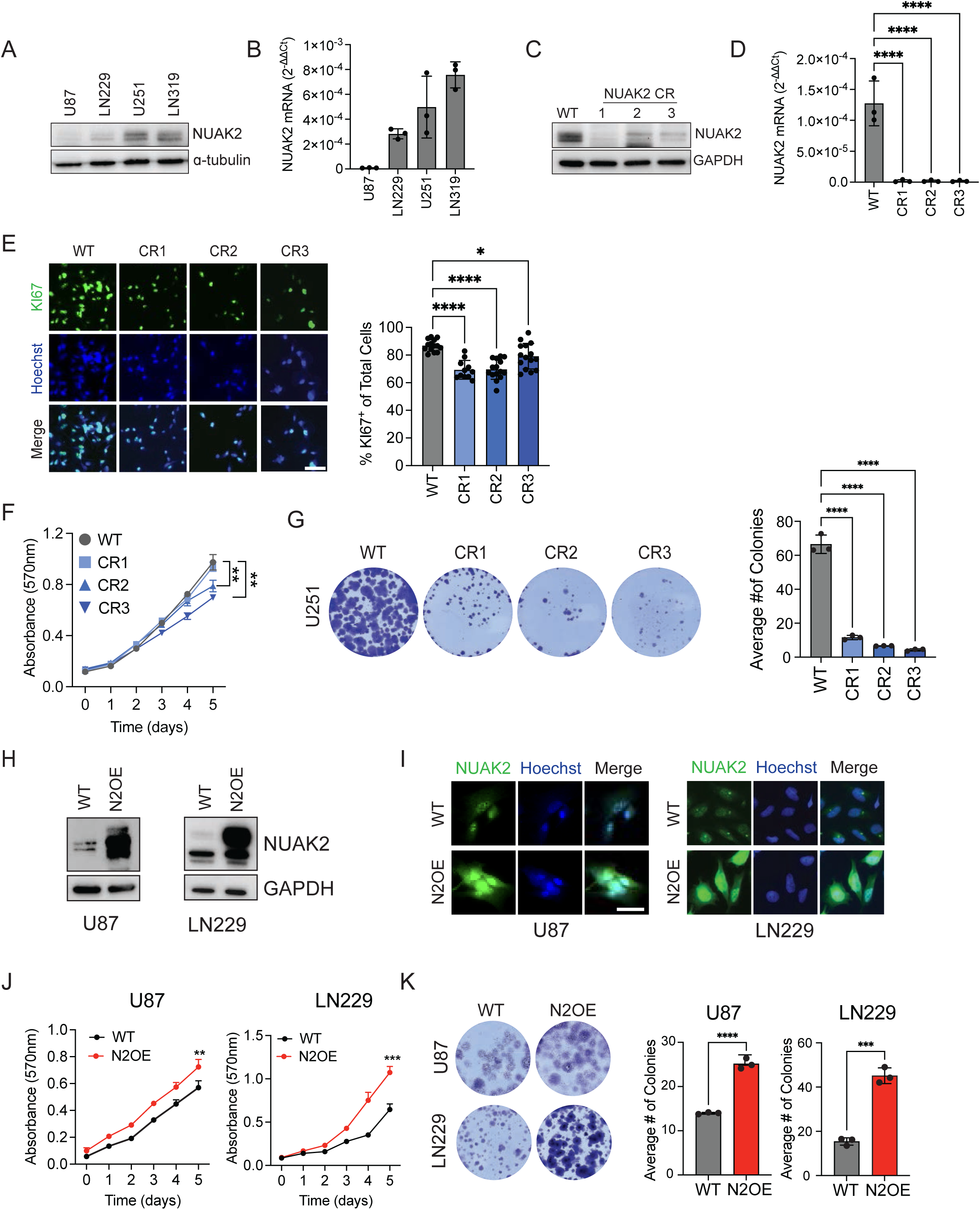
Modulation of NUAK2 expression level is critical for GBM cell proliferation. **A** Representative western blot of NUAK2 protein expression in U87, LN229, U251, and LN319 glioblastoma cell lines. Alpha-tubulin was used as the loading control. **B** qRT-PCR analysis of the mRNA levels of NUAK2 in U87, LN229, U251, and LN319 glioma cell lines. GAPDH was used for normalization. Data are represented as mean ± SD (n=3). **C** Western blot analysis of the efficiency of CRISPR-mediated deletion of NUAK2 in U251 cells. GAPDH was used as a loading control. Three independent clones are shown. Wildtype = WT, NUAK2 CRISPR clone 1 = CR1, CRISPR2 clone = CR2, and CRISPR3 clone = CR3. **D** qRT-PCR analysis of the efficiency of CRISPR-mediated deletion of NUAK2 in U251 cells. Three independent clones are shown. Data are represented as mean ± SD (****p < 0.0001; Statistical significance was determined by one-way ANOVA analysis followed by Dunnett’s multiple comparison test). **E** Representative images of the proliferation marker Ki67 in NUAK2 deleted U251 cells, Hoechst was used to identify cellular nuclei. Quantification of Ki67 is shown on the right (*p = 0.0159, ****p < 0.0001; Statistical significance was determined by one-way ANOVA analysis followed by Dunnett’s multiple comparison test). Scale bar = 50µm. **F** MTT assay evaluating proliferation, as indicated by absorbance, when NUAK2 was deleted in U251 cells. Statistics were evaluated at day 5. WT vs. CR2, WT vs. CR3. Data are represented as mean ± SD (**p < 0.01; Statistical significance was determined by two-way RM ANOVA analysis followed by Uncorrected Fisher’s LSD. Exact p value is reported in Appendix Table S3). **G** Colony formation assay on WT and NUAK2-deleted U251 cells. Quantification of the average number of colonies per well. Data are represented as mean ± SD (n= 3, ****p < 0.001; Statistical significance was determined by one-way ANOVA analysis followed by Dunnett’s multiple comparison test). **H-I** Western blot and immunocytochemical validation of NUAK2 overexpression (N2OE) in U87 and LN229 WT and Nuak2 overexpression (N2OE) cells. NUAK2 is in green, and Hoescht is in blue. GAPDH was used as a loading control. Scale bar = 50µm. **J** MTT assays evaluating the effects of N2OE in U87 and LN229 cells. Data are represented as mean ± SD (**p = 0.0068, ***p = 0.0001; Statistical significance was determined by two-way RM ANOVA analysis followed by Uncorrected Fisher’s LSD). **K** Colony formation assay evaluating the effects of N2OE in U87 and LN229 cells. Data are represented as mean ± SD (***p = 0.0002, ***p <0.001; Statistical significance was determined by unpaired t-test (two-tailed). Exact p value is reported in Appendex Table S3).

Conversely, we conducted NUAK2 overexpression through lentiviral transduction in the two cell lines with the lowest NUAK2 expression, U87 and LN229, to investigate whether upregulated NUAK2 could accelerate GBM cell proliferation. We created NUAK2- overexpressing (N2OE) stable U87 and LN229 cell lines, confirming each cell line’s status with immunoblotting and immunocytochemistry (Fig 2H and I). MTT and colony formation assays revealed that overexpressing NUAK2 significantly enhanced cell proliferation (Fig 2J and K).

Collectively, these findings indicate that NUAK2 is critical for GBM cell growth.

### Silencing NUAK2 impedes GBM cell growth in orthotopic xenograft models

We next evaluated the effect of NUAK2 deletion *in vivo.* We employed a mouse xenograft GBM model in which U251 NUAK2-WT and -CR cells were intracranially injected into BALB/c nude mice. Tumor formation and growth were monitored weekly with *in vivo* bioluminescence imaging (IVIS) from day 7 to day 28 post-injection. The results show significantly smaller tumors in NUAK2-CR mice compared to controls (Fig 3A and B). Histological analysis further confirmed that tumor sizes were markedly smaller in the NUAK2-CR group, with notably fewer Ki67-expressing cells (Fig 3C and D). These findings suggest that NUAK2 deletion effectively suppresses stable tumor engraftment and expansion in the context of the brain.

**Figure 3.**
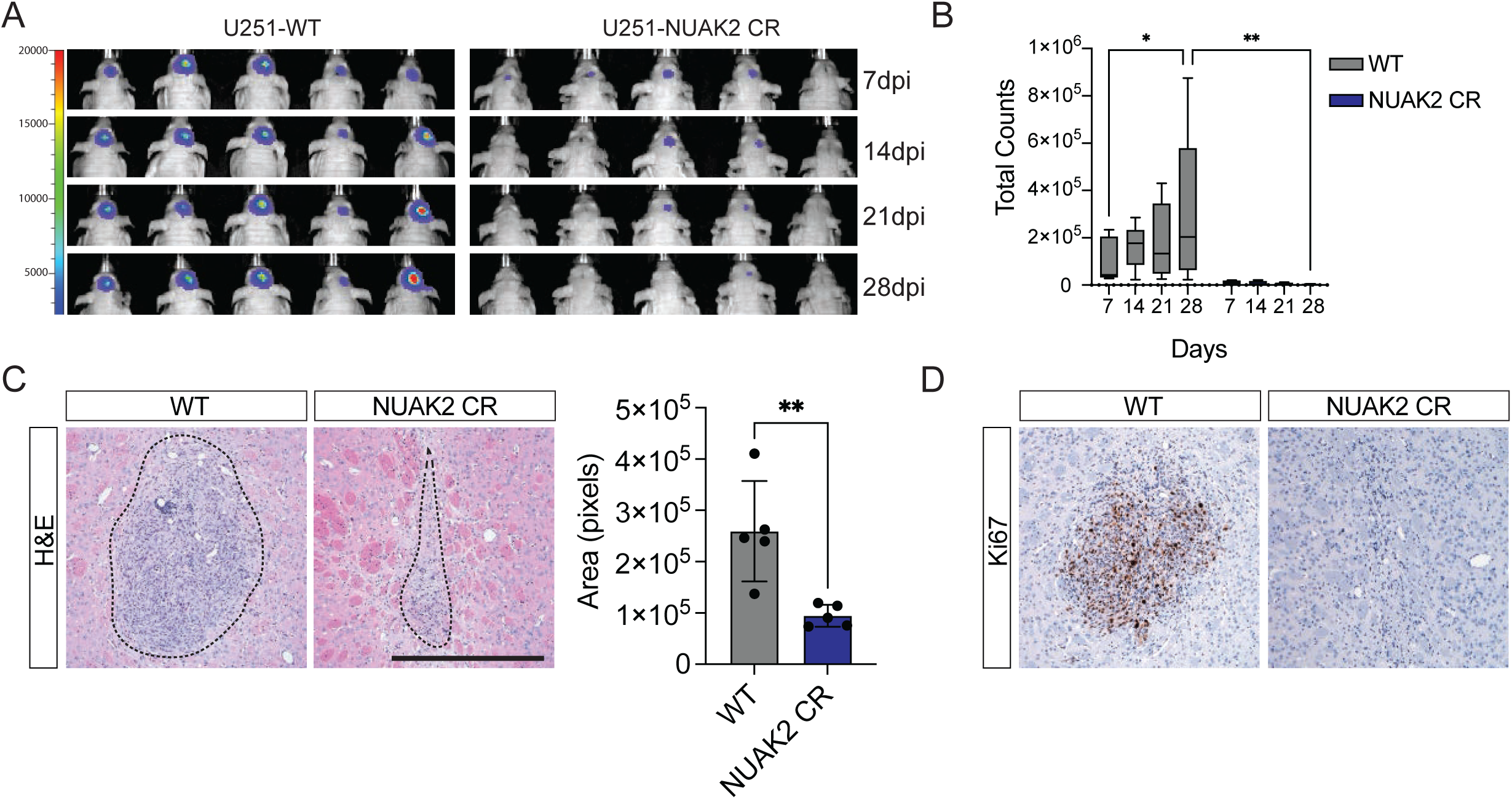
NUAK2 deletion inhibited tumor growth in *in vivo* orthotopic xenografts **A** Intracranial orthotopic xenograft in BALB/c nude Mice using U251 cells with NUAK2 deletion U251-NUAK2 CR) or wildtype (WT) controls (n = 5). Representative bioluminescent images of tumors at 7, 14, 21, and 28 days post-injection are shown. Fluorescence signal intensity is indicated on the left. **B** Quantification of bioluminescent images obtained on IVIS Spectrum imager. Data are represented as mean ± SD (n = 5, *p = 0.036, **p = 0.0044; Statistical significance was determined by two-way RM ANOVA analysis followed by Uncorrected Fisher’s LSD). **C** Representative images of the end-stage tumor (28 dpi) showing H&E staining. Quantification of the area of tumor mass is shown. Data are represented as mean ± SD (n = 5; **p = 0.0062; Statistical significance was determined by unpaired t-test (two-tailed)). Scale bar =500µm. **D** Representative images of the end-stage tumor (28 dpi) showing Ki67 positive proliferating cells.

### *NUAK2 deletion* in a IUE model of malignant glioma supports a role for NUAK2 in GBM

Given the limitation that BALB/c nude mice are immunocompromised, we further examined the role of NUAK2 using a piggyBac *in utero* electroporation (PB-IUE) model of malignant glioma (Chen & LoTurco, 2012; Glasgow et al., 2014; Zhang & Bordey, 2023). PB-IUE-generated tumors are generated in immunocompetent mice and more closely mimic GBM pathophysiology. Using this system, we conducted both NUAK2 GOF and LOF studies to determine the role of NUAK2 in glioma formation.

For NUAK2 LOF (NUAK2-CR) studies, we used a CRISPR-Cas9 approach where dual guide RNAs targeting NUAK2 were co-electroporated with tumor-generating PB-IUE plasmids (Fig 4A). Western blot analysis from harvested tumors confirmed deletion of NUAK2 in electroporated tumors (Fig 4B). Survival studies revealed that the NUAK2-CR cohort had significantly prolonged 50% survival rates compared to control tumor-bearing mice (Fig. 4C). Complementary, GOF studies where tumor-generating PB-IUE constructs were co- electroporated with NUAK2 plasmid, demonstrated that overexpression of NUAK2 (NUAK2- OE) conferred significantly reduced survival rates compared to control mice (Fig 4D). NUAK2 expression in GOF and LOF tumors was validated by immunohistochemical (IHC) analysis (Fig 4E-G; Fig EV1A and B). Proliferation in tumors was analyzed by Ki-67 expression levels revealing that tumors with NUAK2 LOF had fewer proliferating cells, while NUAK2 GOF led to enhanced proliferation (Fig 4E-G; Fig EV1A and B). Notably, NUAK2-OE tumors had large areas of necrosis as compared to control and NUAK2-CR tumors (Fig 4E), consistent with NUAK2-OE leading to excessive growth of the cells and poor prognosis. These results are also consistent with survival trends observed in human glioma patients (Fig 1). Together with our results from orthotopic U251 transplants (Fig 3), these findings indicate that NUAK2 promotes tumorigenesis in our *in vivo* IUE models of high-grade glioma.

**Figure 4.**
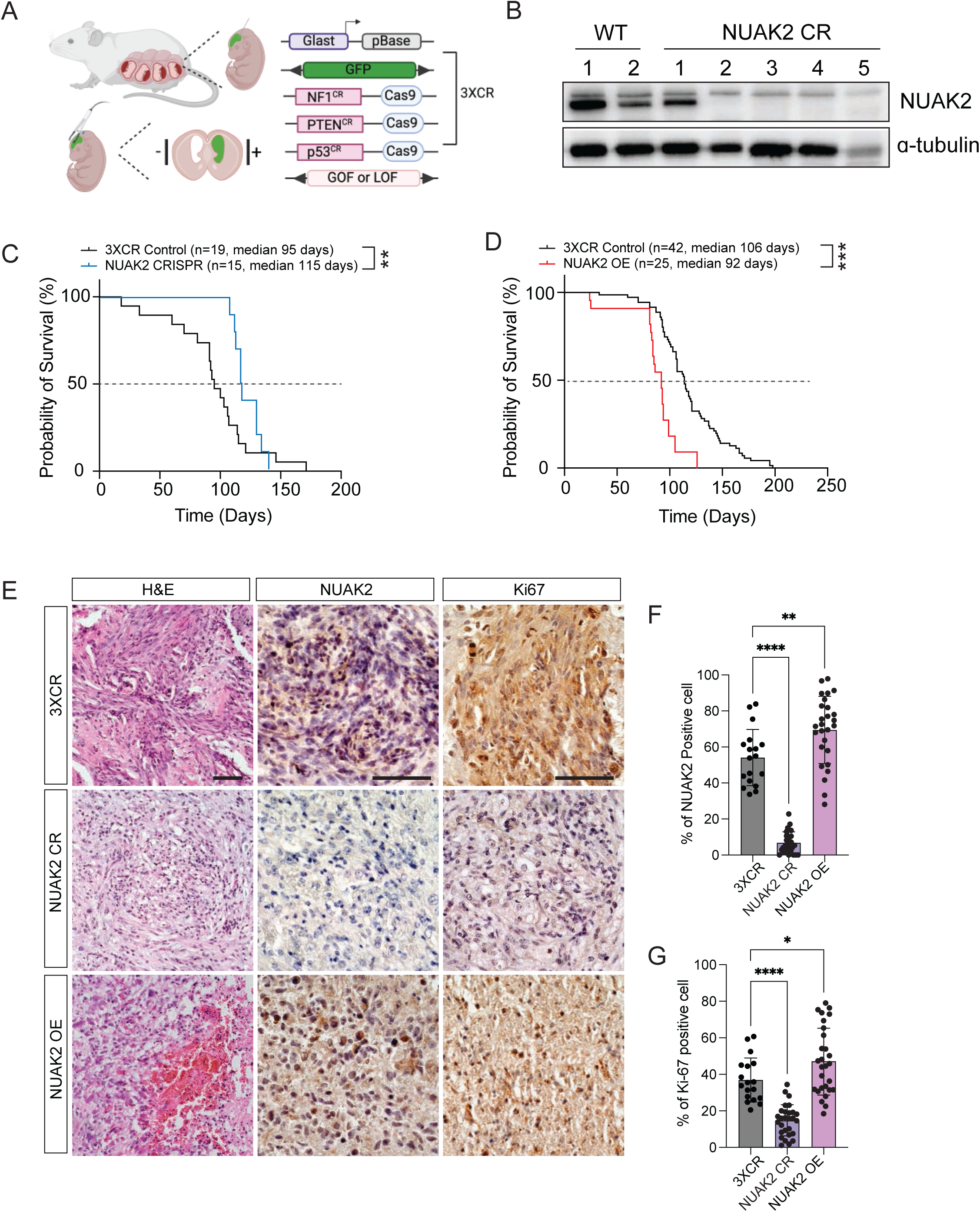
Modulation of NUAK2 expression in an immunocompetent model of malignant glioma affects tumor growth **A** Cartoon representation of the *in utero* electroporation (IUE) model of malignant glioma. **B** Representative western blot of the efficiency of CRISPR-mediated deletion of NUAK2 in IUE-generated malignant gliomas. Alpha-tubulin was used as a loading control. **C** Kaplan-Meier survival analysis from NUAK2 loss-of-function (CRISPR) (n=15) and control tumor-bearing mice (n = 19; **p = 0.001; Statistical significance was determined by log-rank (Mantel-Cox) test). **D** Kaplan-Meier survival analysis from NUAK2 gain-of-function (OE) (n=25) and control tumor- bearing mice (n = 42; ***p = 0.004; Statistical significance was determined by log-rank (Mantel- Cox) test). **E** Representative images of H&E, NUAK2 expressing, and Ki67 positive proliferating cells in control, NUAK2 deleted or OE tumors at P50. Scale bar =100µm. **F** Quantification of NUAK2 expression in control, NUAK2 deleted or OE tumors. Data are represented as mean ± SD (**p = 0.0015, ****p < 0.0001; Statistical significance was determined by one-way ANOVA analysis followed by Dunnett’s multiple comparison test). **G** Quantification of Ki67 positive proliferating cells in control, NUAK2 deleted or OE tumors. Data are represented as mean ± SD (*p = 0.0282, ****p < 0.0001; Statistical significance was determined by one-way ANOVA analysis followed by Dunnett’s multiple comparison test).

### NUAK2 mediates mesenchymal transition through ECM regulation

To investigate the mechanistic role of NUAK2 in GBM, we performed bulk RNA-seq transcriptomic analysis using NUAK2-WT and NUAK2-CR U251 cell lines. Deletion of NUAK2 resulted in 658 (273 upregulated and 385 downregulated; Dataset EV1) differentially expressed genes (DEGs) with a log2 fold change greater than two in U251-CR cells compared to control cells (Fig EV2A). Gene ontology (GO) analysis of the biological process and cellular component groups using NUAK2-CR DEGs highlighted top annotations related to the extracellular matrix (ECM; Fig 5A; Dataset EV2). To validate these findings, we conducted GO analysis using RNA-seq data from TCGA GBM patient samples sourced from the GlioVis data portal. This analysis suggests that ECM-related terms are significantly influenced by NUAK2 expression in our samples (Fig 5B), demonstrating consistent results not only in homogeneous cell populations but also in actual GBM patient samples.

**Figure 5.**
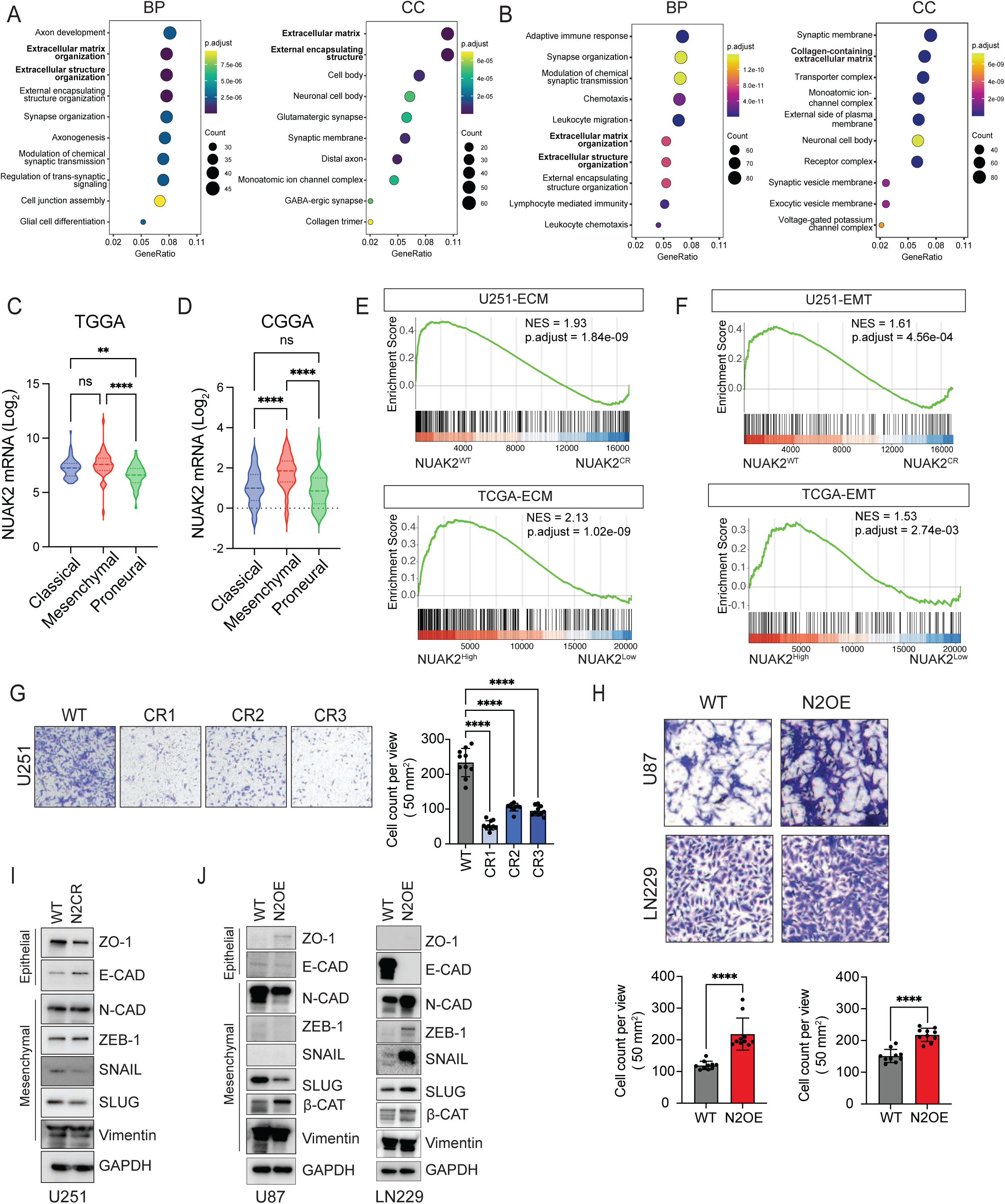
NUAK2 mediates mesenchymal transition through ECM regulation **A** Gene Ontology (GO) term enrichment analysis of U251 DEGs after NUAK2-CRISPR mediated deletion. Plots represent DEG categories by Biological Process (BP) and Cellular Component (CC). **B** Gene Ontology (GO) term enrichment analysis of TCGA DEGs of NUAK2^Low^ Group compared to NUAK2^High^. Plots represent DEG categories by Biological Process (BP) and Cellular Component (CC). **C** Distribution of Log2 value of NUAK2 mRNA expression in GBM subtypes based on TCGA. Violin-plot shows a significant association between proneural vs. classical and proneural vs. mesenchymal subtypes. Data are represented as mean ± SD (**p = 0.0042, ****p < 0.0001; Statistical significance was determined by one-way ANOVA analysis followed by Tukey’s multiple comparison test). **D** Distribution of Log2 value of NUAK2 mRNA expression in GBM subtypes based on CCGA. Violin-plot shows a significant association between classical vs. mesenchymal and mesenchymal vs. proneural subtypes. Data are represented as mean ± SD (****p < 0.0001; Statistical significance was determined by one-way ANOVA analysis followed by Tukey’s multiple comparison test). **E-F** GSEA of U251 and TCGA-GBM DEGs enrichment in extracellular matrix (ECM) and epithelial-to-mesenchymal transition (EMT) gene sets. **G** Representative images of transwell assay migration assay post NUAK2 deletion in U251 cells in three independent CRISPR clones. Quantification is shown on the right panel as mean ± SD (n = 10, ****p < 0.0001; Statistical significance was determined by one-way ANOVA analysis followed by Dunnett’s multiple comparison test). **H** Representative images and quantification of transwell assay after NUAK2 overexpression in U87 and LN229 cells. Data are represented as mean ± SD (n = 10; ****p < 0.0001; Statistical significance is determined by unpaired t-test (two-tailed)). **I** Representative western blot images of EMT markers from U251 NUAK2 deleted (N2CR) lysates. Gapdh was used as a loading control. **J** Representative western blot images of EMT markers from U87 and LN229 NUAK2 overexpressing lysates. Gapdh was used as a loading control.

Since ECM regulation is closely linked to epithelial-to-mesenchymal transition (EMT) in GBM (Khoonkari et al., 2022; Majc et al., 2020; Mohiuddin & Wakimoto, 2021; So et al., 2021), we hypothesized that NUAK2 may play a pivotal role in mesenchymal GBM and mediate EMT through ECM modulation. To investigate this, we analyzed TCGA GBM RNA-seq data from the GlioVis data portal to evaluate NUAK2 expression across GBM subtypes, finding significantly higher NUAK2 levels in the mesenchymal group compared to the proneural subtype, which is the least aggressive GBM phenotype (Fig 5C and D). Gene Set Enrichment Analysis (GSEA) further supported these findings, showing enrichment of mesenchymal signatures in NUAK2 ^WT^ and NUAK2^High^ samples, while NUAK2 ^CR^ and NUAK2^Low^ samples were enriched in proneural signatures (Fig EV2B; Table EV3). These findings suggest that NUAK2 loss leads to reduced mesenchymal properties.

Next, we examined the relationship between ECM modulation and the EMT process. Both ECM signature and EMT-related genes were positively enriched in NUAK2-WT U251 cells and NUAK2^High^ GBM patients, suggesting that ECM regulation and EMT are interdependent processes (Fig 5E and F). A mesenchymal signature is related to more migratory properties, therefore we examined whether cellular migration is affected by U251 expression using transwell assays; finding that NUAK2 loss impairs cell migration, while NUAK2 overexpression enhances migration (Fig 5G and H). Additionally, immunoblotting from NUAK2-OE or NUAK2- CR cell lysates revealed decreased epithelial markers and increased mesenchymal markers with higher NUAK2 levels compared to controls (Fig 5I and J), further supporting the role of NUAK2 in promoting EMT via ECM modulation.

To identify critical regulators of ECM modulation, we identified the set of overlapping DEGs from NUAK2-CR U251 cells and NUAK2^Low^ GBM patients in the ECM signature. Eleven genes were consistently altered across both datasets (Fig EV2C). qRT-PCR validation confirmed that these ECM genes are closely linked to NUAK2-driven GBM progression and expansion (Fig EV2D). Taken together, these results indicate that NUAK2 promotes EMT and facilitates GBM progression through its regulation of ECM-related genes.

### The NUAK2 inhibitor HTH-02-006 attenuates GBM cell proliferation

After identifying the critical role of NUAK2 expression in glioblastoma cell progression, we investigated whether inhibition of NUAK2 kinase activity could mimic the effects of NUAK2 gene depletion in GBM cells. We utilized a commercially available NUAK2 inhibitor, HTH-02- 006 (Fu et al., 2022; Yuan et al., 2018), across four GBM cell lines.

Since HTH-02-006 is a semi-specific NUAK2 inhibitor that could potentially inhibit NUAK1, its homolog, we investigated the potential relevance of NUAK1 to GBM. Our analysis revealed that NUAK1 expression is relatively stable throughout brain development in both mice and humans (Fig EV3A-D), distinct from the tightly controlled regulation of NUAK2. This suggests that NUAK1 expression is not developmentally dynamic across stages. Additionally, NUAK1 mRNA levels were significantly lower in both low- and high-grade gliomas than in normal brain tissues (Fig EV3E). Further examination of glioblastoma subtypes showed no significant differences in NUAK1 expression across subtypes (Fig EV3F and G), indicating a lack of strong correlation with tumor grade. Survival analyses from Kaplan-Meier curves also showed no significant relationship between NUAK1 expression and glioma patient prognosis (Fig EV3H and I). These findings collectively suggest that NUAK1 has limited relevance in glioblastoma tumorigenesis and progression, indicating that the effect of HTH-02-006 is more likely mediated through NUAK2.

To determine the effect of HTH-02-006 in GBM cells, we performed MTT and colony formation assays. We observed that the inhibitor suppressed cell proliferation in a dose-dependent manner across all four cell lines. However, sensitivity to the drug varied depending on the level of NUAK2 expression in the cell line (Fig 6A and B; Fig EV4A and B). Notably, U87 cells showed a limited response to the inhibitor, likely due to their lower NUAK2 expression levels.

**Figure 6.**
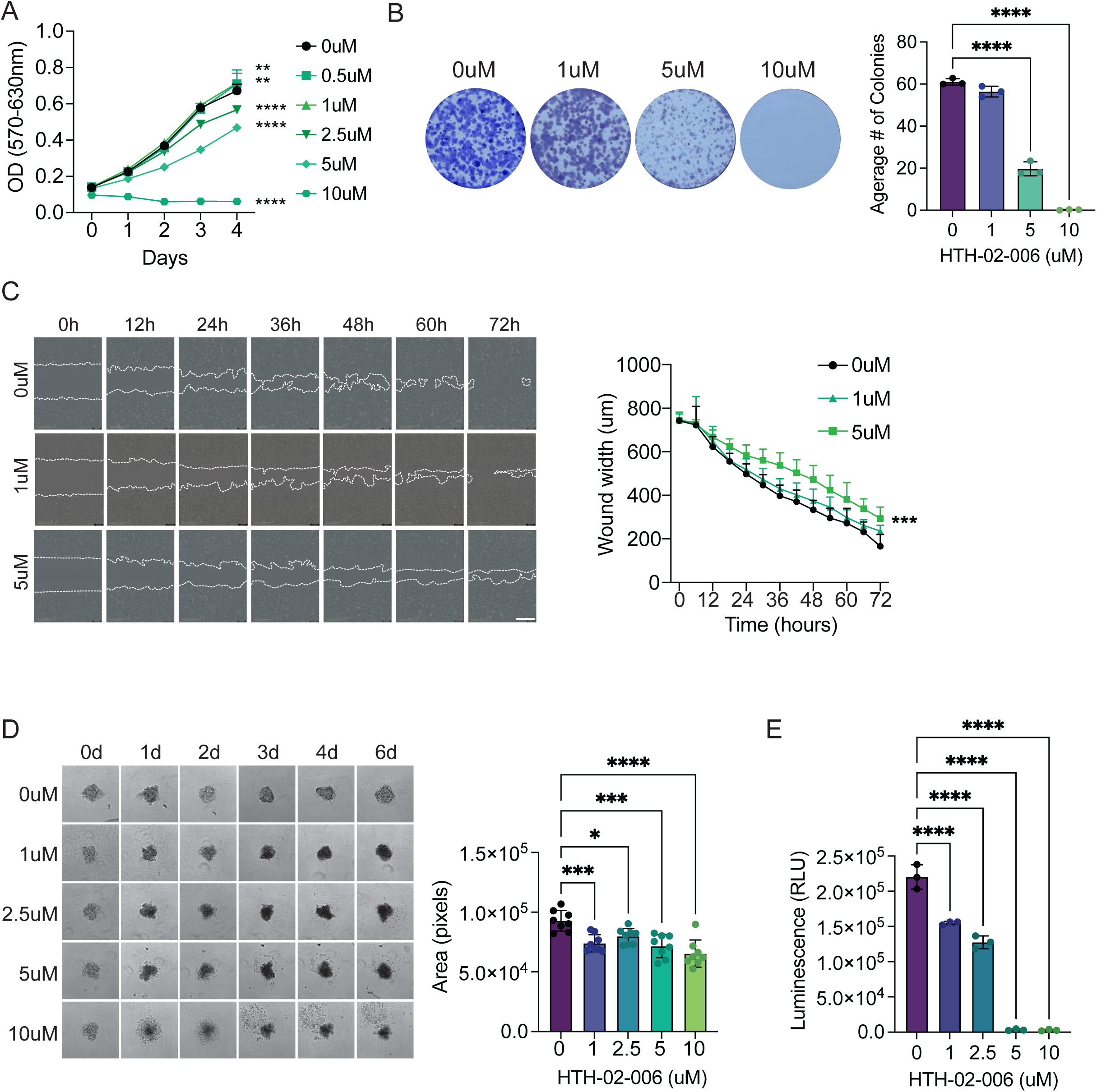
Pharmaceutical inhibition of NUAK2 suppresses GBM cell progression and expansion. **A** MTT assay for proliferation in HTH-02-006 treated U251 cells. Data are represented as mean ± SD (n = 7; **p < 0.001, ****p < 0.0001; Statistical significance was determined by two- way RM ANOVA analysis followed by Uncorrected Fisher’s LSD. Exact p values are reported in Appendix Table S3). **B** Colony formation assay and quantification on HTH-02-006 treated U251 cells. Data are represented as mean ± SD (n= 3; ****p < 0.0001; Statistical significance was determined by one-way ANOVA analysis followed by Dunnett’s multiple comparison test). **C** Representative phase images and quantification of HTH-02-006 treated U251 cell migration into the wound area. Data are represented as mean ± SD (n= 7; ***p = 0.0008; Statistical significance was determined by two-way RM ANOVA analysis followed by Uncorrected Fisher’s LSD). White dotted lines demarcate the wound boundary. **D** Representative brightfield images and quantification of total spheroid area of HTH-02-006 treated U251 spheroids over the course of 6 days. Data are represented as mean ± SD (n= 8; *p < 0.05, ***p < 0.001, ****p < 0.0001; Statistical significance was determined by one-way ANOVA analysis followed by Dunnett’s multiple comparison test. Exact p values are reported in Appendix Table S3). **E** Cell Titer Glo 3D cell viability assay quantification in HTH-02-06 treated U251 spheroids at day 6. Data are represented as mean ± SD (n= 3; ****p < 0.0001; Statistical significance was determined by one-way ANOVA analysis followed by Tukey’s multiple comparison test).

This finding suggests that the growth and propagation of U87 cells may be less dependent on NUAK2 activity. Furthermore, scratch wound healing analysis showed that the inhibitor markedly hindered the migration of GBM cells (Fig 6C; Fig EV4D). However, clear migration activity was not observed in U87 cells, likely due to their distinct growth pattern, characterized by convergence and the formation of circular clusters.

While our analysis using two-dimensional (2D) monolayer cultures provides initial insights into the inhibitor efficacy, this method does not replicate the architecture of tumor masses *in vivo*. To address this issue, we employed three-dimensional (3D) spheroid analysis to evaluate the effects of HTH-02-006 on glioblastoma cells. The 3D spheroid model is particularly advantageous when drug kinetics are not well understood in organisms, as it incorporates *in vivo*-like features such as cell-cell interactions, drug penetration, and ECM deposition (Barbosa et al., 2021; Zanoni et al., 2016). HTH-02-006 treatment demonstrated dose-dependent growth inhibition in all four GBM spheroid models, underscoring its clinical significance (Fig 6D and E; Fig EV5). Collectively, these findings support the conclusion that NUAK2 is a promising therapeutic target for GBM.

## DISCUSSION

A growing number of studies indicate that cancer cells capitalize on embryonic developmental paradigms to promote their development and progression. The parallels between development and cancer most commonly relate to stemness, EMT, and proliferation which give the cell a selective growth advantage (Cao et al., 2023; Sharma et al., 2022). Indeed, cell proliferation and migration are fundamental processes in both normal development and cancer. However, proliferation in development follows tightly regulated pathways, while cancers exploit these mechanisms for uncontrolled growth (Aiello & Stanger, 2016, 2016; Balachandran & Narendran, 2023; Ma et al., 2010). Therapies targeting abnormal developmental pathways in cancer have been developed, but are limited by the need to identify specific actionable targets (Dempke et al., 2017; Kiesslich et al., 2012). Therefore, investigating the factors that intersect embryogenesis and tumorigenesis is critical for understanding tumor biology and developing more effective therapeutic strategies. In this study, we identified NUAK2 as a fetal oncogene essential for CNS development which also plays a pivotal role in glioma tumorigenesis and progression. Importantly, pharmaceutical suppression of NUAK2 can attenuate these processes.

NUAK2 is highly expressed in a range of cancers, including melanoma, prostate, and hepatic tumors. (W. Fu et al., 2022; Namiki et al., 2011; Yuan et al., 2018). In these tumors, NUAK2 has been shown to have tumor-promoting properties, facilitating proliferation, migration, and invasion of these cancer cells. In GBM, NUAK2 expression has been reported to be high in glioblastoma tissue compared to adjacent normal brains, linking NUAK2 to glioblastoma tumorigenesis. Fu *et al*. identified that the microRNA miR-143 inhibits glioblastoma progression, in part by degrading NUAK2 (Fu et al., 2016). However, the relationship between NUAK2 expression in normal developing and adult brain tissue and its tumorigenic role in the CNS remains unclear. In this study, we demonstrate that while NUAK2 is essential for CNS development, its aberrant expression in adult brains contributes to tumorigenesis by mimicking developmental processes such as proliferation and migration.

Our *in vitro* and *in vivo* LOF and GOF studies of NUAK2 determined that it can regulate GBM progression. Moreover, our transcriptomic analysis of GBM cells and TGCA patient data that express high or low levels of NUAK2 revealed that NUAK2 likely exerts its effects by modulating the ECM. ECM has emerged as a critical factor driving malignancies, including gliomas (Huang et al., 2021; Larriba et al., 2024; Venning et al., 2015; Zhao et al., 2021). In gliomas, ECM supports tumor progression by facilitating cell invasion and proliferation, particularly via epithelial-mesenchymal transition (Khoonkari et al., 2022; Majc et al., 2020; Mohiuddin & Wakimoto, 2021; So et al., 2021). It plays a significant role in promoting resistance to therapy by modulating tumor density and stiffness, making drug treatments or radiation difficult to penetrate the tumor and reach the proliferating inner tumor mass.

Furthermore, ECM remodeling alters tissue stiffness, activating pathways that drive tumor growth, making it a potential therapeutic target (Mohiuddin & Wakimoto, 2021; Wei et al., 2024). Despite these findings, the mechanisms by which NUAK2 is reactivated and drives tumorigenesis are not fully elucidated, and further comprehensive and interdisciplinary investigations need to be done.

Interestingly, there have been multiple reports about NUAK2 and cellular stresses. NUAK2 has been linked to cellular responses to metabolic stress, such as glucose/glutamine deprivation, UVB exposure, and treatments with AICAR and metformin (Lefebvre et al., 2001; Lefebvre & Rosen, 2005). Cells are continually subjected to mechanical and chemical stresses, which can result in accumulated mutations and, ultimately, uncontrolled growth and migration—hallmarks of cancer. Therefore, it is possible that NUAK2 dysregulation in the brain may occur as a stress response, but further studies are required to connect NUAK2, cellular stress, and glioma tumorigenic processes more definitively.

To the best of our knowledge, this study is the first to investigate the effects of both genetic and pharmaceutical modulation of NUAK2 on GBM cells, proposing a novel approach for GBM-specific precision treatment. We observed the growth of four glioblastoma cell lines was effectively blocked by HTH-02-006 treatment both in attached and suspension culture. HTH-02-006 is a NUAK2 semi-specific inhibitor. However, it has good NUAK2 selectivity with limited off-target effects (Yuan et al., 2018). While HTH-02-006 is roughly nine times more specific to NUAK2 than to its closely related homolog NUAK1, it is important to consider the NUAK1 in glioma cells due to its substantial role in the brain (J. Courchet et al., 2013; V. Courchet et al., 2018; Lanfranchi et al., 2024; Lasagna-Reeves et al., 2016). Therefore, we assessed the relevance of NUAK1 in glioma patients by analyzing its expression patterns in comparison to normal brain tissue and evaluating survival rates based on NUAK1 expression levels. We found that NUAK1 did not share the same dynamic embryonic expression pattern as NUAK2, nor did its expression correlate with glioma tumor grade or patient survival. This is in contrast to a recent finding that NUAK1 was correlated to patient survival and had a role in promoting GBM growth (Lu et al., 2013). The discrepancies between these findings and our observations likely arise from differences in the study populations, which differ in demographics, environmental and/or clinical contexts. To further validate the drug efficacy in GBM, we need to better understand the specific contexts in which NUAK2 operates in glioblastoma and to understand how the downstream effects of NUAK2 kinase activity regulate major signaling pathways such as HIPPO, WNT, TGFβ, and others known to influence glioma tumorigenesis. Importantly, further efforts to develop more specific, effective, and especially brain-bioavailable inhibitors for clinical application are needed.

## MATERIALS AND METHODS

### Animals

CD-1 IGS (Charles River #022) timed pregnant mice were used for IUE studies. BALB/c nude immunocompromised mice were obtained from the University of California San Diego (UCSD) in-house breeding program for xenograft studies. Care of all animals in this study was approved by the UCSD Institutional Animal Care and Use Committee (IACUC) and followed NIH guidelines and procedures.

### Cell Culture and Reagents

Four GBM cell lines (U87MG, LN229, U251MG, LN319) were used in this study. U87MG and LN229 glioblastoma cell lines were purchased from ATCC (#HTB-14, #CRL-2611). U251MG and LN319 cell lines were obtained from Addexbio (#C0005029, #C0005001). All cells were maintained in humidified incubators at 37℃ and 5% CO2. Cell lines were tested for mycoplasma using the LookOut Mycoplasma PCR Detection Kit (Sigma; MP0035). U87 was grown in Dulbecco’s Modified Eagle Medium/F12 (DMEM/F12) supplemented with 10% fetal bovine serum (FBS) and 1% penicillin-streptomycin (PS). LN229, U251MG, and LN319 were cultured in DMEM with 10% FBS and 1% PS. HTH-02-006 (#AOB36960) was purchased from Aobious and dissolved in dimethyl sulfoxide (DMSO)

### Orthotopic Xenograft Models

7-8 week-old male BALB/c nude mice were used to generate cell line xenograft models. U251 wildtype and CRISPR-edited cells were dissociated with trypsin and resuspended at PBS (1.7 x 10^5^ cells/ul). 5 x 10^5^ cells in 3ul of PBS were injected into the specific coordinates (x, y, z = 0.5, -2.0, -3.0) from bregma using stereotaxic injection system (RWD Life Science).

Bioluminescence-based *in vivo* imaging of xenograft mice was performed at 7, 14, 21, and 28 days after cell injection using a Perkin Elmer IVIS Spectrum imaging system. Mice were intraperitoneally injected with 10μL/g body weight of 15mg/ml D-luciferin, anesthetized, and placed in IVIS Spectrum bioluminescent and fluorescent imaging systems (Perkin Elmer).

Luminescence signals were developed and acquired per minute followed by one minute exposure time for 10-15 minutes. To quantify bioluminescent intensity a region of interest (ROI) was selected and analyzed using IVIS software. Brains were harvested, fixed, and embedded for histology analysis on the 28th-day post-injection.

### *In Utero* Electroporation (IUE)

*In utero* electroporation was used to generate mouse gliomas as previously described (Chen & LoTurco, 2012; Glasgow et al., 2017). In short, uterine horns of E15 pregnant females were exposed and the appropriate DNA cocktail containing 1X Fast Green dye indicator was injected into the lateral ventricles of embryos. The embryos were then electroporated with BTW Tweezertrodes connected to a BTX 8300 electroporator. The settings for electroporation were: 33V, 55ms per pulse conducted six times, at 100ms intervals. DNA combinations used were the helper plasmid pGLAST-PBase (2.0 μg/μL) pbCAG-GFP, pbCAG-Luciferase, crNF1, crPTEN, and crp53, and either NUAK2-expressing or NAUK2-targeting sgRNA plasmids, all at a concentration of 1.0 μg/μL each (Chen & LoTurco, 2012; Glasgow et al., 2017; John Lin et al., 2017). Mouse specific NUAK2 sgRNAs (Yuan et al., 2018) targeting exon 1 (5’- CCTCGCGGTCCCCGCACCAT-3’ and 5’-CTACGAGTTCCTGGAGACGC-3’) and non-targeting control (5’-ATGTTGCAGTTCGGCTCGAT-3’) were cloned into pX330. Animals were sacrificed at various time points and processed for further analysis.

### Stable Cell Line Generation

To generate NUAK2 knockout lines in human glioblastoma cell lines, CRISPR guides targeting human NUAK2 (5’-TGGAGTCGCTGGTTTTCGCG-3’) were cloned into a GFP- or mCherry- containing lentiviral vectors: LentiCRISPRv2GFP (Addgene#82416) and LentiCRISPRv2mCherry (Addgene #99154), respectively. Hek293T cells were transfected with the NUAK2 sgRNA expressing GFP and mCherry vectors and the appropriate viral packaging plasmids using Viafect Transfection Reagent (Promega #E981) according to the manufacturer’s instructions. The virus was collected over three days, combined, and filtered prior to the transduction of human GBM cell lines. Transduced cells expressing both GFP and mCherry were enriched using Fluorescence-activated Cell Sorting (FACS). After transduction, cells were trypsinized and resuspended in 1 ml of FACS sorting buffer (0.1% BSA, 1% pen/strep, 1% 1 M HEPES pH 7, 25 mg/ml DNase in Leibowitz medium (Fisher, #21083027). Green/Red double-positive cells were sorted into 96-well plates which was performed by UC San Diego Human Embryonic Stem Cell Core Facility using a BD FACSAriaII. Clones were cultured in a 96-well until 80%-90% confluency, then transferred to plates with larger surfaces. From a 24-well plate, clones were screened by PCR and propagated for further experimentation.

### Cell proliferation and Clonogenic assays

MTT assays were conducted for two-dimensional proliferation assays using an MTT assay kit (Roche; #11465007001) following the manufacturer’s protocol. 1X10^3^ cells per well were plated in 96 well plates. To validate the effect of the NUAK2 inhibitor, cells were treated with a complete growth medium containing various concentrations of HTH-02-006 (1, 2.5, 5, 10, 20uM) for the indicated amount of time. 0.1% dimethyl sulfoxide (DMSO) was used as a control. Upon collection, HTH-02-006 treated cells were labeled with 10ul of labeling solution per well for four hours and lysed by adding 100ul of solubilization reagent followed by overnight incubation. The 570 and 630nm absorbance were measured using a spectrophotometer (Perkin Elmer).

For colony formation assay, 500 cells/well were plated in 6 well plates and maintained for 10- 14 days. To evaluate the effect of HTH-02-006, 1000 cells/well were plated in 6 well plates and grown for a week, then the cells were treated with various concentrations of the inhibitor for another week. Next, the colonies were fixed in 100% methanol and stained with 0.05% crystal violet solution. Excessive stains were removed by rinsing the plates with tap water. The plates were air-dried and photographed for quantification.

### Scratch wound healing assay

Cells were seeded at 2X10^4^ cells per well in 96 well plates. After 24 hours, uniform wounds were created using IncuCyte 96-well WoundMaker Tool as described in the manufacturer’s protocol. After the scratch wound creation, cells were carefully washed twice with 1X PBS, treated, and maintained at indicated concentrations of HTH-02-006. The wound closure process was visualized every 12 hours for three days and analyzed in real-time with the IncuCyte S3 live-cell imaging system (Sartorius Bioscience).

### Transwell migration assay

Cells were grown in regular media to 60% confluency in 10 cm plates. On the next day, the media was changed to serum-free media to starve the cells overnight. Then, 500ul of complete media including FBS, was placed into the wells of a 24-well-plate. Transwell inserts (Thermo Fisher; # 07-200-150) were transferred into each well, creating an upper chamber. Serum- starved cells were harvested and 1.5X10^4^ cells/well were resuspended in 400µL of serum-free medium and plated onto the upper chamber of the transwell insert. Cells were allowed to migrate while incubating at 37°C for 40-46 hours. Next, media was gently removed from the inserts and washed with PBS, followed by fixation with 800µL of 4% paraformaldehyde (PFA) in PBS split between the lower and upper chambers. After 15 minutes of fixation at room temperature inserts were washed twice with PBS. Cells were permeabilized with 100% cold methanol for 10 min, washed twice with PBS, and stained with 0.05% crystal violet for 15 min. Inserts were washed twice with PBS and non-migrated cells were removed by gently scraping with cotton swabs. Membranes were then cut out, fixed in permount on a slide, and imaged on an Olympus BX63 Microscope.

### 3D spheroid analysis

To generate 3D spheroids of each GBM cell line, 1X10^3^ cells suspended in serum-free media were plated in each well of ultra-low attachment 96 well plates (Corning; #07-201-680) and briefly spun down by centrifugation. Spheroids were treated with various concentrations of HTH-02-006 after they formed circular masses, then imaged daily until day 6 after the initial drug treatment with ImageXpress MicroXLS (Molecular Devices) from the UCSD screening core laboratory. To determine viable cells in the spheroids, the Cell Titer Glo 3D Cell Viability Assay reagent (Promega; #G9681) was used as described in the manufacturer’s protocol.

To determine spheroid sizes, batch image analysis of spheroids was conducted in Fiji (version 2.16.0) using a script to measure spheroid area. External plugins used in the script are AdjustableWatershed, BioVoxxel (version 2.6.0), and MorpholibJ (version 1.64). Spheroid image annotations were manually inspected for quality. Valid spheroid area measurements accurately traced the perimeter of the spheroid while excluding the surrounding cell monolayer. Dissociated spheroids were counted as having an area of zero. Out of the 1,152 spheroid images for the four cell lines (U87, U251, LN229, and LN319), 36 images were manually annotated. For manual annotation, the area of the dissociated spheroids was set to zero, or the spheroid was manually traced in yellow, and its contained area was measured. Four images were removed due to poor quality.

### Quantitative Real-Time PCR (qRT-PCR)

Total RNA was isolated using Trizol (Invitrogen) solution following the manufacturer’s protocol. Trizol reagent was added directly to cells or tissues and lysates were either immediately processed or stored at -80℃. The RNA concentrations were measured with a NanoDrop spectrophotometer (Thermo Fisher). cDNAs were generated from 0.5ug of total RNA per sample by reverse transcription using iScript cDNA synthesis kit (Biorad; #1708891). Samples were analyzed by CFX384 real-time system (Biorad) using PerfeCTa® SYBR® Green FastMix® (Quantabio; #101414-270) according to the manufacturer’s protocol. Gene expression was normalized to a housekeeping gene *GAPDH.* See Appendix Table S1 for the list of qPCR primers used in the study.

### Western Blot

Cells were lysed in radioimmunoprecipitation assay (RIPA; 10 mM Sodium chloride, 50 mM Tris-HCl, 1% NP-40, 0.5% Sodium deoxycholate, 0.1% SDS) buffer with ethylenediaminetetraacetic acid (EDTA) free protease/phosphatase inhibitor cocktail (Thermo Fisher; #78441) and kept at -20°C for long-term storage and future analysis. Protein concentration was determined by performing Bradford assay (Sigma; #B6916). A total of 20- 40ug of protein lysates were resolved with polyacrylamide gel electrophoresis (8-10% Tris-HCl SDS PAGE gels) and transferred onto either nitrocellulose or polyvinylidene difluoride (PVDF) membrane based on the molecular weight of the target protein. Membranes were then immersed in 3% bovine serum albumin (BSA) and incubated for one hour at room temperature followed by overnight incubation with primary antibodies at 4℃. Membranes were washed with Tris-buffered saline (TBS) with 0.1% Tween and then incubated with secondary antibodies for one hour. Lastly, the target protein signal was developed using Western Blotting Luminol reagent (Santacruz Biotechnology; #sc-2048). See Appendix table S2 for primary antibodies and specifications used for the study.

### Immunohistochemistry (IHC-P)

For paraffin embedding, mice were perfused with PBS followed by 10% neutral buffered formalin for whole-body fixation. Fixed brains were dissected and drop-fixed in 10% neutral buffered formalin for 16 hours followed by 24 hours incubation in 70% ethanol. The brains were processed for paraffin embedding at the UCSD Biorepository and Tissue Technology core. Brains were sectioned at 5um using a Leica microtome and allowed to dry for analysis. Sections were deparaffinized using xylene and a series of decreasing ethanol concentration washes. Sections were washed with TBS-T and antigen retrieval was performed using sodium citrate buffer (pH 6.0) at 95℃ for 15 minutes. Immunohistochemistry was performed using ImmPRESS® Excel Amplified Polymer Staining Kit (Vector Laboratories; #MP-7601). Briefly, sections were washed with TBS-T before using BLOXALL® Endogenous HRP/AP Blocking Solution for 10 minutes followed by two washes with TBS-T. Sections were blocked in 2.5% Horse serum for 30 minutes followed by incubation in primary antibodies, either Ki-67 (Cell Signaling Technologies; D3B5) 1:500 or NUAK2 (Novus Biologicals; NBP1-81880) 1:50 antibodies overnight at 4℃. Sections were washed three times with TBS-T before applying Amplifier Antibody (Goat Anti-Rabbit IgG) for 15 minutes. Sections were washed three times with TBS-T and ImmPRESS Horse Anti-Goat IgG Polymer Reagent was applied for 30 minutes. Before chromogenic detection, sections were washed two times with TBST before the DAB substrate was applied and allowed to develop for two minutes. The slides were washed three times with TBS-T before counterstaining with hematoxylin and dehydrating through a series of increasing ethanol concentrations and xylene incubations. Sections were mounted and dried for 24 hours prior to imaging. The percentage of positively stained cells was analyzed using QuPath software cell detection protocol on 20X images of tumor areas. Three separate 500x500 pixel squares were counted for each sample.

### Immunocytochemistry (ICC)

Circular glass coverslips (Fisher; #50949008, 12mm) were coated with 0.01% poly-L-ornithine solution (Sigma; P4957) overnight prior to cell seeding. On the next day, an appropriate number of cells were plated onto the coverslips to yield ∼70% confluency. After treatment cells were fixed using cold 4% PFA for 15 minutes followed by permeabilization with 0.1% triton X- 100 in PBS for three minutes with gentle agitation. The coverslips were washed with PBS and incubated with 3% BSA blocking buffer (3% BSA in PBS (w/v)) for one hour at room temperature. After blocking, primary antibodies diluted in the same blocking buffer were added onto coverslips and incubated overnight at 4℃, Coverslips were then washed with PBS and incubated with fluorescence-conjugated secondary antibodies for 1-2h at room temperature, washed with PBS, and nuclei stained with Hoechst 33258 (Sigma; #B2883). Coverslips were mounted using an anti-fade mounting medium (Vectashield; H-1400), dried overnight, and imaged using the fluorescence microscope (Olympus). Images were analyzed and quantified using FIJI software.

### RNA-sequencing (RNA-seq)

#### Sample Preparation

Total RNA was isolated using TRIzol Reagent following the manufacturer’s protocol. The quality of total RNA was evaluated using an Agilent Tapestation 4200, and only samples with an RNA Integrity Number (RIN) above 9.0 were selected for RNA-seq library preparation with the Illumina® Stranded mRNA Prep kit (Illumina, San Diego, CA). Library preparation was conducted according to the manufacturer’s protocol by the UCSD Institute for Genomic Medicine (IGM) Core Facility. The prepared libraries were multiplexed and sequenced using 100 base pair (bp) paired-end reads (PE100) on an Illumina NovaSeq 6000, achieving a sequencing depth of approximately 25 million reads per sample. Demultiplexing was performed with the bcl2fastq Conversion Software (Illumina, San Diego, CA).

### Data Analysis

For U251 RNA-seq analysis, FASTQ files were processed in Galaxy using Trimmomatic with default parameters. Read alignment was performed with HISAT2 using the hg38 reference genome and default parameters. Raw expression data was obtained using featureCounts with default parameters. Differential expression analysis was performed in R (version 4.4.1) using the DESeq2 package (version 1.46.0). Differentially expressed genes (DEGs) were determined based on a significance threshold adjusted p-value of < 0.05 and a log2 fold change (LFC) > 2. 685 DEGs, with 273 upregulated and 385 downregulated, were identified. Differential expression analysis for TCGA GBM was obtained from the open-access web application for data visualization and analysis, GlioVis, which compared the highest 25% and lowest 25% NUAK2 expressing samples. DEGs were determined based on a significance threshold adjusted p-value < 0.05 and an LFC > 1. 1494 DEGs, with 807 upregulated and 687 downregulated, were identified.

### Over-Representation Analysis

Over-representation analysis (ORA) was performed using the clusterProfiler package (version 4.14.0). Gene Ontology (GO) terms for Biological Process (BP) and Cellular Component (CC) categories were identified using the set of 17,767 genes for U251 and 20,501 for TCGA GBM as background. A significance threshold of p < 0.01 and q < 0.05 was applied. The ‘simplify’ method from clusterProfiler with default parameters was used to remove redundant GO terms. The Benjamini-Hochberg procedure was used for multiple- hypothesis testing correction.

### Gene Set Enrichment Analysis

Gene Set Enrichment Analysis (GSEA) was done using clusterProfiler, with genes ranked by the Wald statistic generated from DESeq2 for U251 and calculated by GlioVis for TCGA_GBM. GO: BP terms were analyzed using default parameters with a minimum gene set size of 50. GSEA using gene sets for the EMT, MES signature, and PN signature were obtained from MSigDb under the systematic names M817, M2122, and M2115, respectively. The EMT gene set is equivalent to the GO Biological Process term “epithelial to mesenchymal transition.” Analysis using these gene sets was performed separately with default parameters. A p-value cutoff of 0.05 was used for all analyses. Benjamini-Hochberg procedure was used for multiple-hypothesis correction.

### Statistical analysis

Statistical analyses were conducted using GraphPad Prism 10 software. The data represent findings from at least three independent experiments. Unpaired t-tests were used to assess significance (p < 0.05). Kaplan–Meier survival curves were generated, and survival comparisons were evaluated using the Log-rank (Mantel-Cox) test. More statistics information is reported in Appendix Table S3.

## ACKNOWLEDGEMENTS

We would like to thank the University of California San Diego (UCSD) IGEM Core, the UCSD Human Embryonic Stem Cell Core Facility (hESCCF), and the Screening Core Laboratory directed by Dr. Jair Siqueira-Neto of the UCSD Center for Drug Discovery Innovation for their expert assistance. Figure 4A was created with BioRender.com.

## CONFLICT OF INTEREST STATEMENT

### Contributions

HJ and SG conceived and directed the project. HJ, AD, SJ, and SG wrote and edited the manuscript. HJ, AD, WY, EM, SJ, and SG performed and analyzed experiments. All authors gave comments and approved the final manuscript.

### Corresponding author

Correspondence to Stacey M. Glasgow.

## ETHICS STATEMENT

Care of all animals in this study followed NIH guidelines and procedures were approved by the UCSD Institutional Animal Care and Use Committee (IACUC). All animal experiments were performed in accordance with the approved protocols and guidelines. Intracranial tumor size was monitored using bioluminescent imaging and animals were sacrificed if they showed signs of distress or pain.

## DATA AVAILABILITY STATEMENT

Data generated for this manuscript will be made available upon reasonable request to the corresponding author. The RNA-sequencing results have been deposited to the Gene Expression Omnibus (GEO) and can be found under the accession number GSE285513.

All scripts and RNA-sequencing analysis code can be found in the following GitHub repository: https://github.com/smglasgowlab/nuak2-2024

## Expanded Figures

**Figure EV 1.**
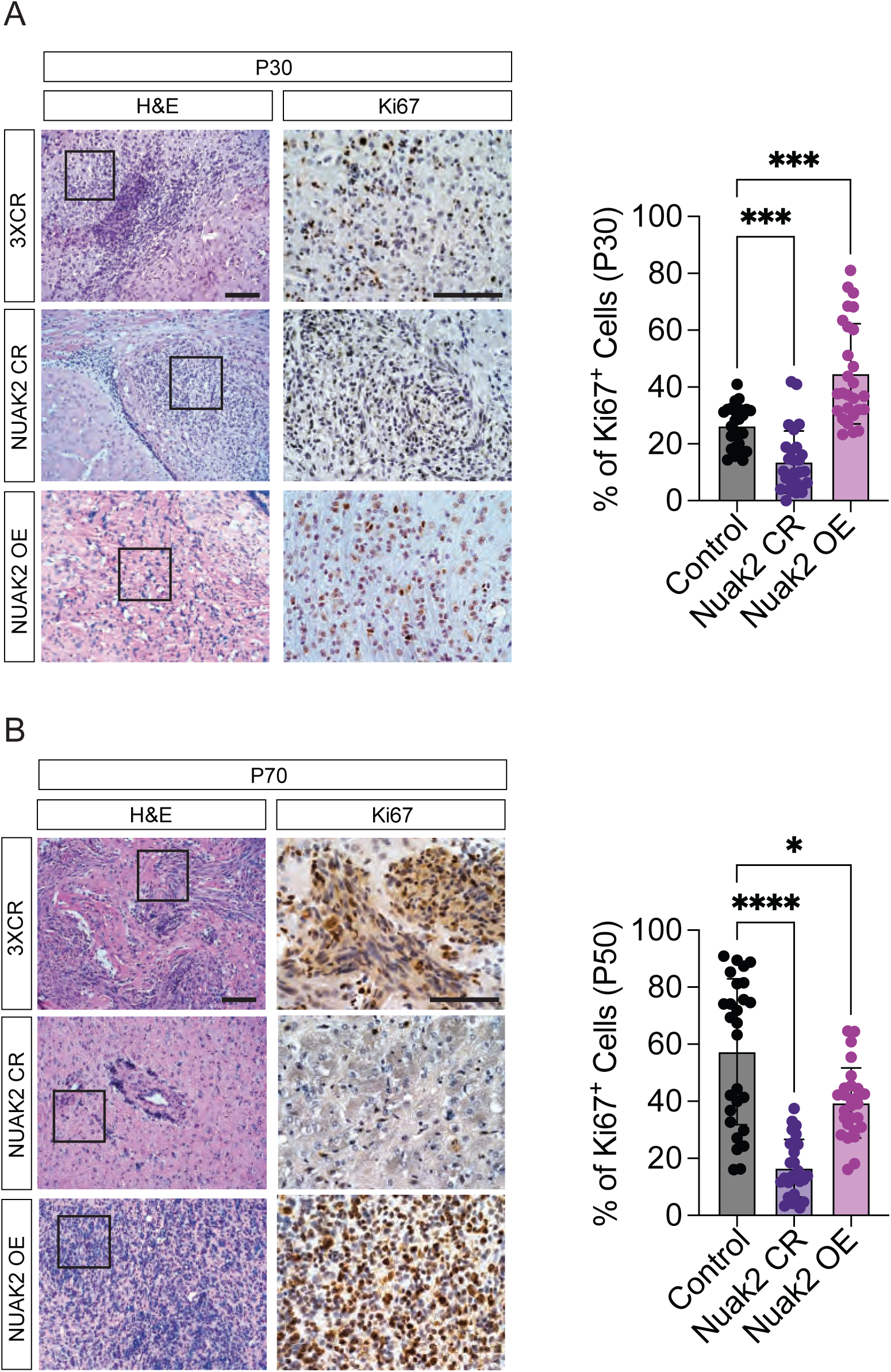
Histological analysis of P30 and P70 IUE. **A** Representative images of H&E and Ki67 proliferating cells in control, NUAK2 deleted or OE tumors at P30. Scale bar = 100µm. Quantification analysis of Ki67 positive cells is represented as mean ± SD (***p = 0.0001; Statistical significance was determined by two-way RM ANOVA analysis followed by Dunett’s multiple comparisons test). **B** Representative images of H&E and Ki67 proliferating cells in control, NUAK2 deleted or OE tumors at P70. Scale bar = 100µm. Quantification analysis of Ki67 positive cells is represented as mean ± SD (*p = 0.016, ****p < 0.001; Statistical significance was determined by two-way RM ANOVA analysis followed by Dunett’s multiple comparisons test).

**Figure EV 2.**
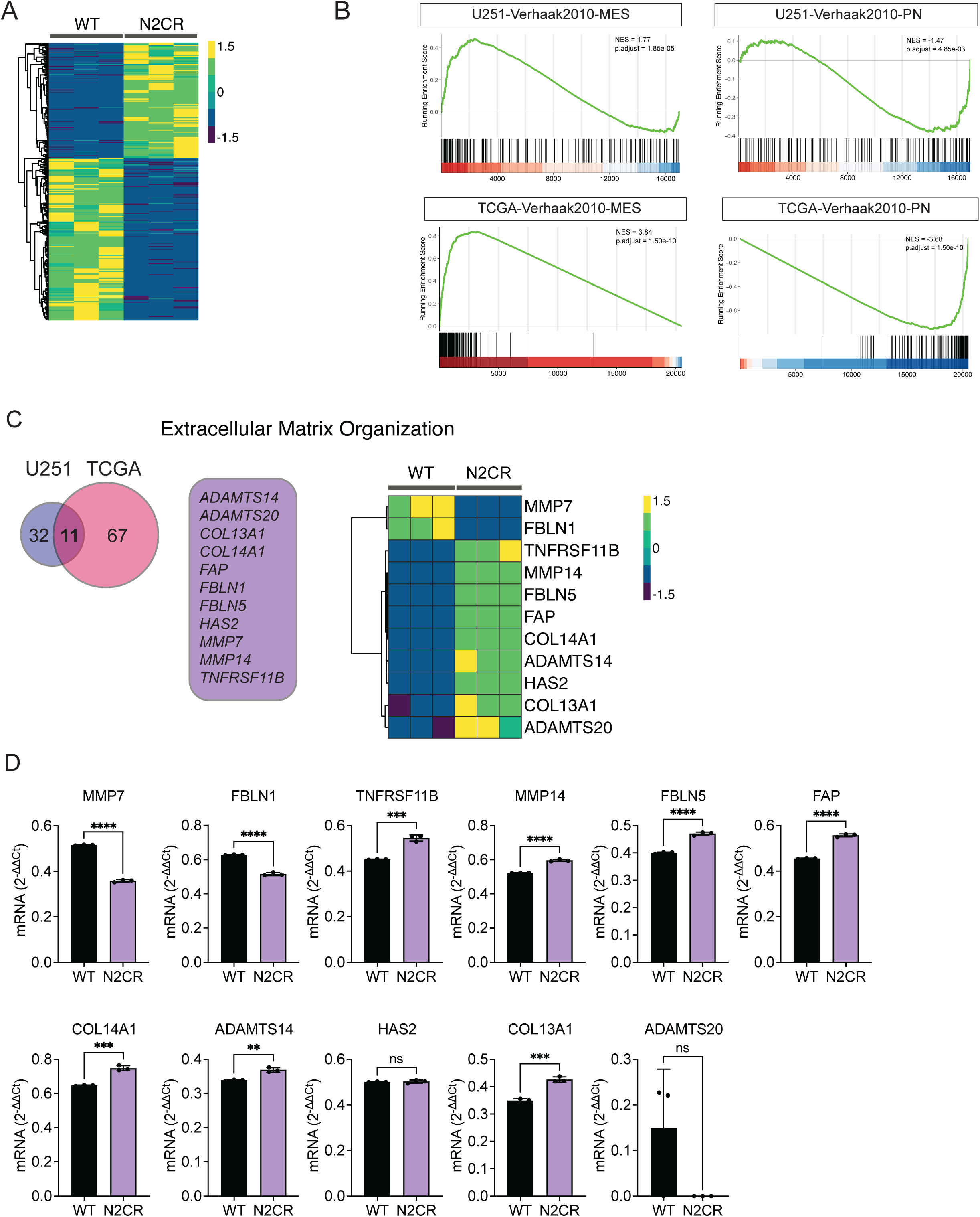
Putative migration regulatory genes identified from NUAK2 ECM associated GO group. **A** Heatmap of differentially expressed genes in U251 control and U251 NUAK2 CRISPR- deleted cells. **B** GSEA enrichment plots of U251 NUAK2-CR and TCGA NUAK2^Low^ gene lists versus queried gene lists from either mesenchymal (MES) or proneural (PN) are shown. **C** Venn Diagram depicting the correlation between NUAK2-deleted U251 cells and TCGA- GBM ECM-associated DEGs. Heatmap of 11 shared genes between U251 NUAK2-deleted cells and TCGA-GBM ECM DEGs. **D** qRT-PCR of the 11 shared ECM genes from control and U251 NUAK2-deleted cells. Data are represented as mean ±SD (n= 3, *p < 0.01, ***p < 0.001, ****p < 0.0001; Statistical significance is determined by unpaired t-test (two-tailed). Exact p values are reported in Appendix Table S3).

**Figure EV 3.**
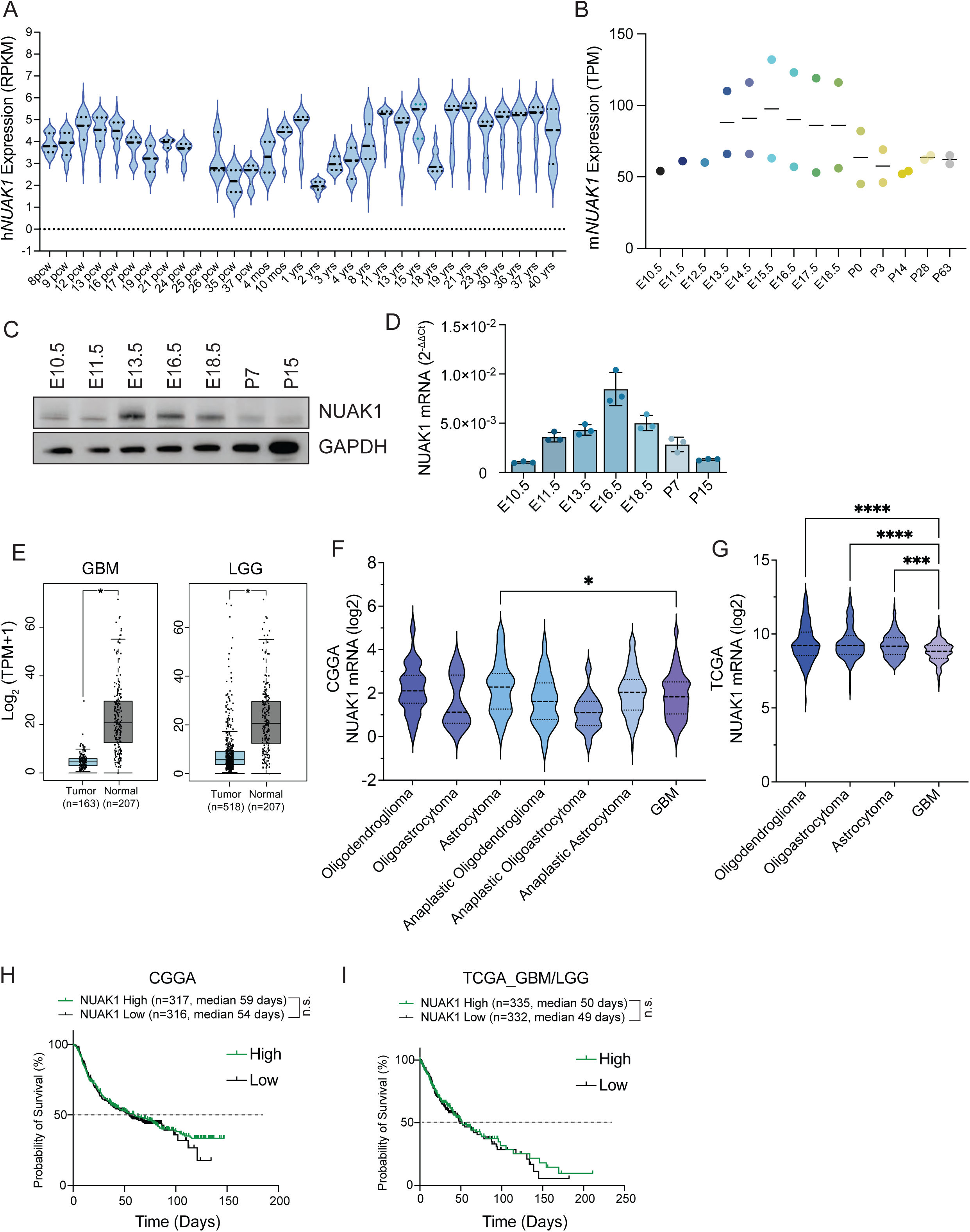
NUAK1 is not associated with GBM progression and patient survival **A** RPKM-normalized NUAK1 mRNA expression of specific human brain regions from 8 post- conception weeks (pcw) to 40 years of age. Data was obtained from the BrainSpan Atlas. **B** TPM-normalized NUAK1 mRNA expression of mouse forebrain or hindbrain ranging from embryonic day 10.5 to postnatal day 63. Data was obtained from EMBL’s European Bioinformatics Institute (EMBL-EBI; https://www.ebi.ac.uk/). **C** Representative western blot of NUAK1 protein expression in wildtype embryonic brain tissue across 7 stages of development. GAPDH was used as the loading control. **D** Representative RT-PCR of NUAK1 mRNA expression in wildtype embryonic brain tissues across developmental stages. **E** Normalized NUAK1 mRNA expression of TCGA GBM (n = 163) or LGG (n = 518) and GTEx non-tumor (n = 207) samples (*p < 0.05; Statistical significance is determined by one-way ANOVA). Data was obtained from the GlioVis Database. **F** NUAK1 mRNA expression across glioma subtypes in the CGGA dataset. Data are represented as mean ±SD (*p = 0.0193; Statistical significance is determined by one-way ANOVA followed by Tukey’s multiple comparisons test). **G** NUAK1 mRNA expression across glioma subtypes in the TCGA dataset. Data are represented as mean ±SD (****p < 0.0001; Statistical significance is determined by one-way ANOVA followed by Tukey’s multiple comparisons test). **H** Kaplan-Meier survival analysis from CGGA of high (21 days; n = 317) and low (145 days; n = 316) NUAK1 expressors shows no correlation with survival outcomes (p = 0.682; Statistical significance was determined by log-rank (Mantel-Cox) test). **I** Kaplan-Meier survival analysis from TCGA of high (15 days; n = 335) and low (134 days; n = 332) NUAK2 expressors shows no correlation with survival outcomes (p = 0.6262; Statistical significance was determined by log-rank (Mantel-Cox) test).

**Figure EV 4.**
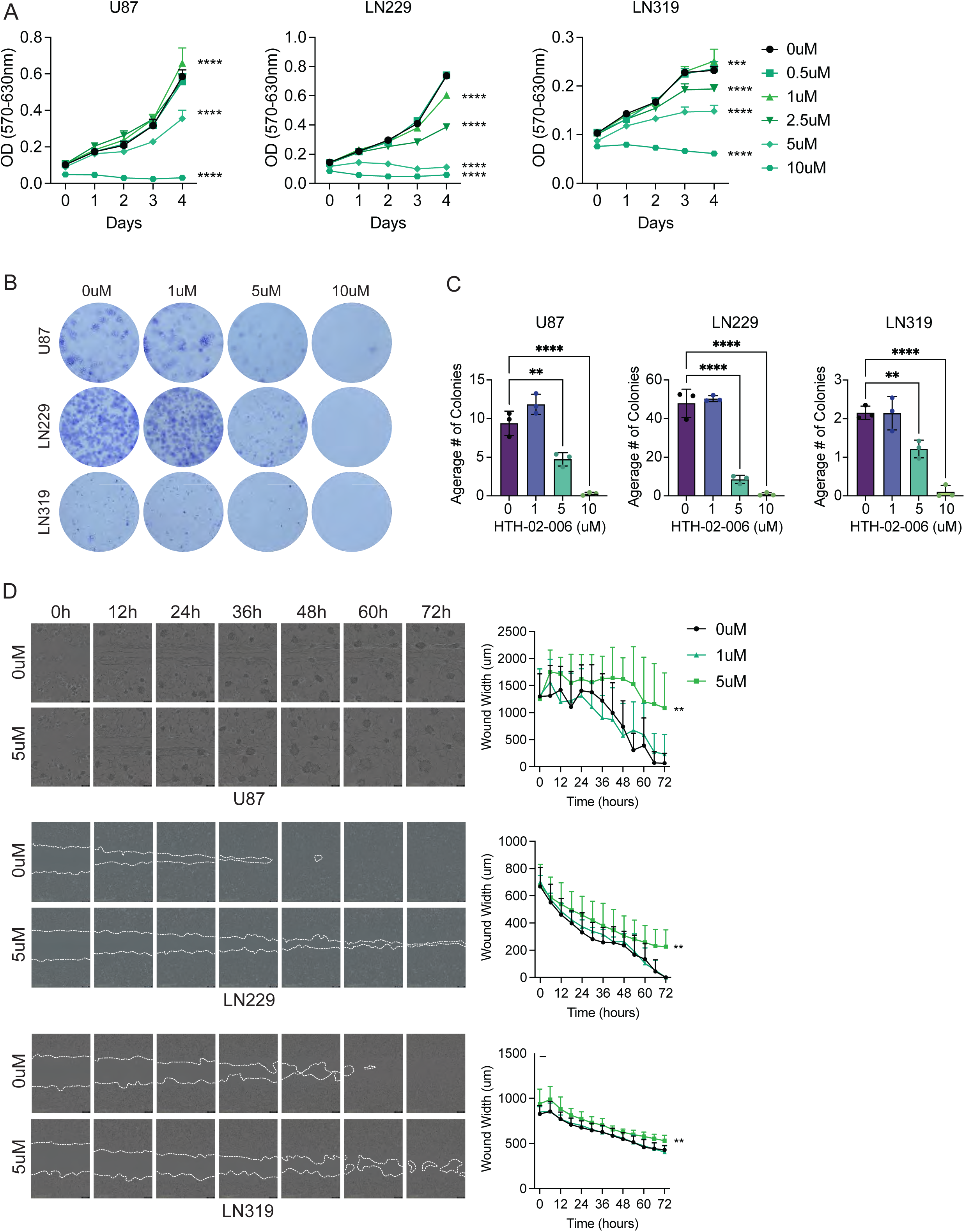
NUAK2 inhibitor, HTH-02-006, attenuates GBM cell progression. **A** MTT assay for proliferation in HTH-02-006 treated U87, LN229, and LN219 cells. Data are represented as mean ±SD (***p = 0.0008, ****p < 0.0001; Statistical significance was determined by two-way RM ANOVA followed by Dunnett’s multiple comparison test. Exact p values are reported in Appendix Table S3). **B** Representative images of colony formation assay of U87, LN229, and LN319 cells with HTH-02-006 treatment. **C** Quantification of colony formation assay (n = 3, **p < 0.01, ****p < 0.0001; Statistical significance was determined by one-way ANOVA followed by Dunnett’s multiple comparison test. Exact p values are reported in Appendix Table S3). **D** Representative images and quantification of HTH-02-06 treated U87, LN229, and LN319 cell migration into the wound area. Data are represented as mean ±SD (**p < 0.01; Statistical significance was determined by two-way RM ANOVA followed by Dunnett’s multiple comparison test. Exact p values are reported in Appendix Table S3). The white dotted lines demarcate the wound boundary.

**Figure EV 5.**
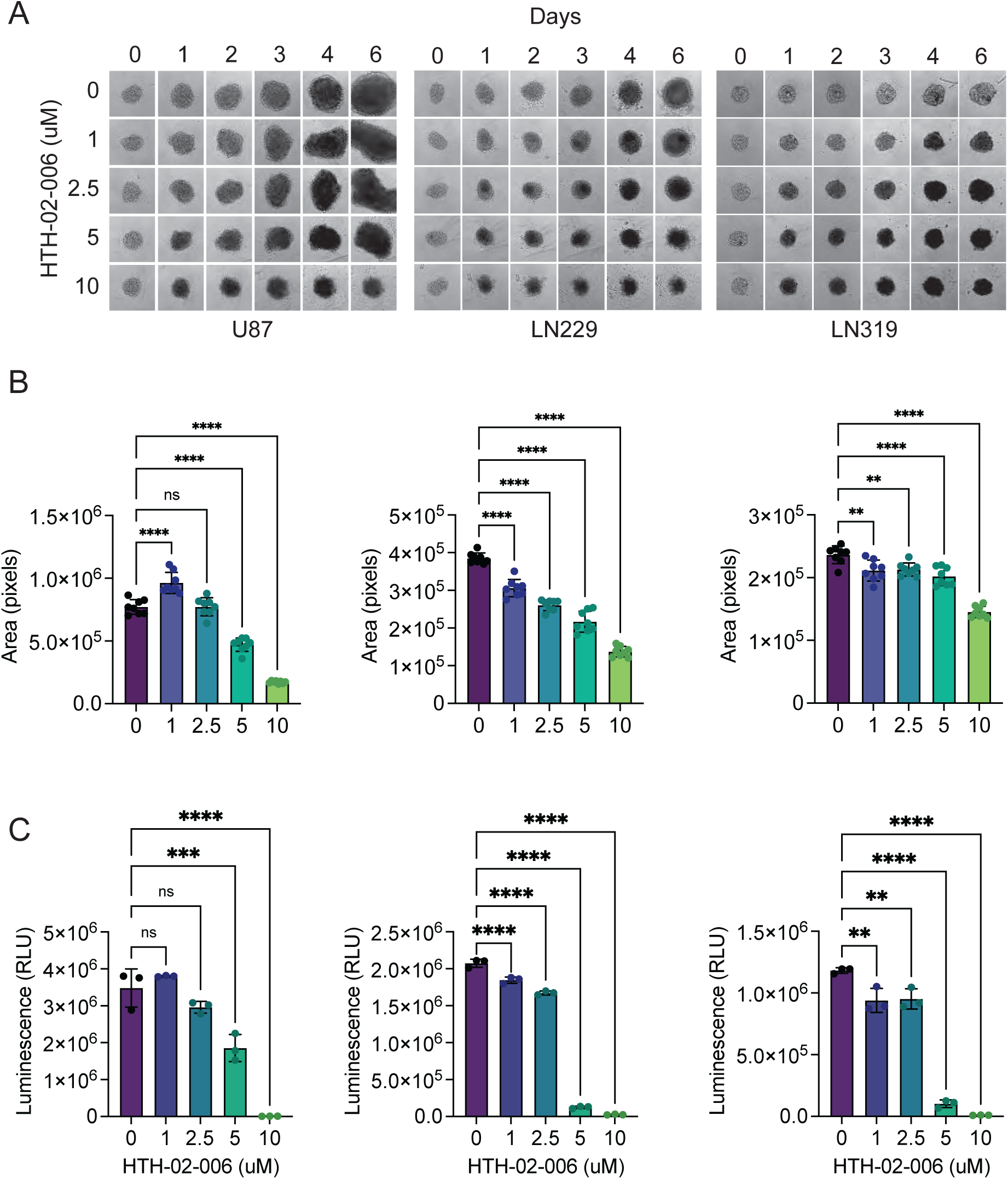
Efficacy of HTH-02-006 in 3D GBM spheroids. **A** Representative brightfield images of spheroid assay in HTH-02-06 treated U87, LN229, and LN319 cells over the course of 6 days. **B** Quantification of total spheroid area of HTH-02-06 treated U87, LN229, and LN319 cells spheroids. Data are represented as mean ±SD (n = 8, **p < 0.01, ****p < 0.0001; Statistical significance was determined by one-way ANOVA followed by Dunnett’s multiple comparison test. Exact p values are reported in Appendix Table S3). **C** Luminescence intensity of viable cells in HTH-02-06 treated U87, LN229, and LN319 spheroids at day 6. Data are represented as mean ±SD (n = 3, **p < 0.01, ***p < 0.001, ****p < 0.0001; Statistical significance was determined by one-way ANOVA followed by Dunnett’s multiple comparison test. Exact p values are reported in Appendix Table S3).

## APPENDIX

**Appendix Table S1.**
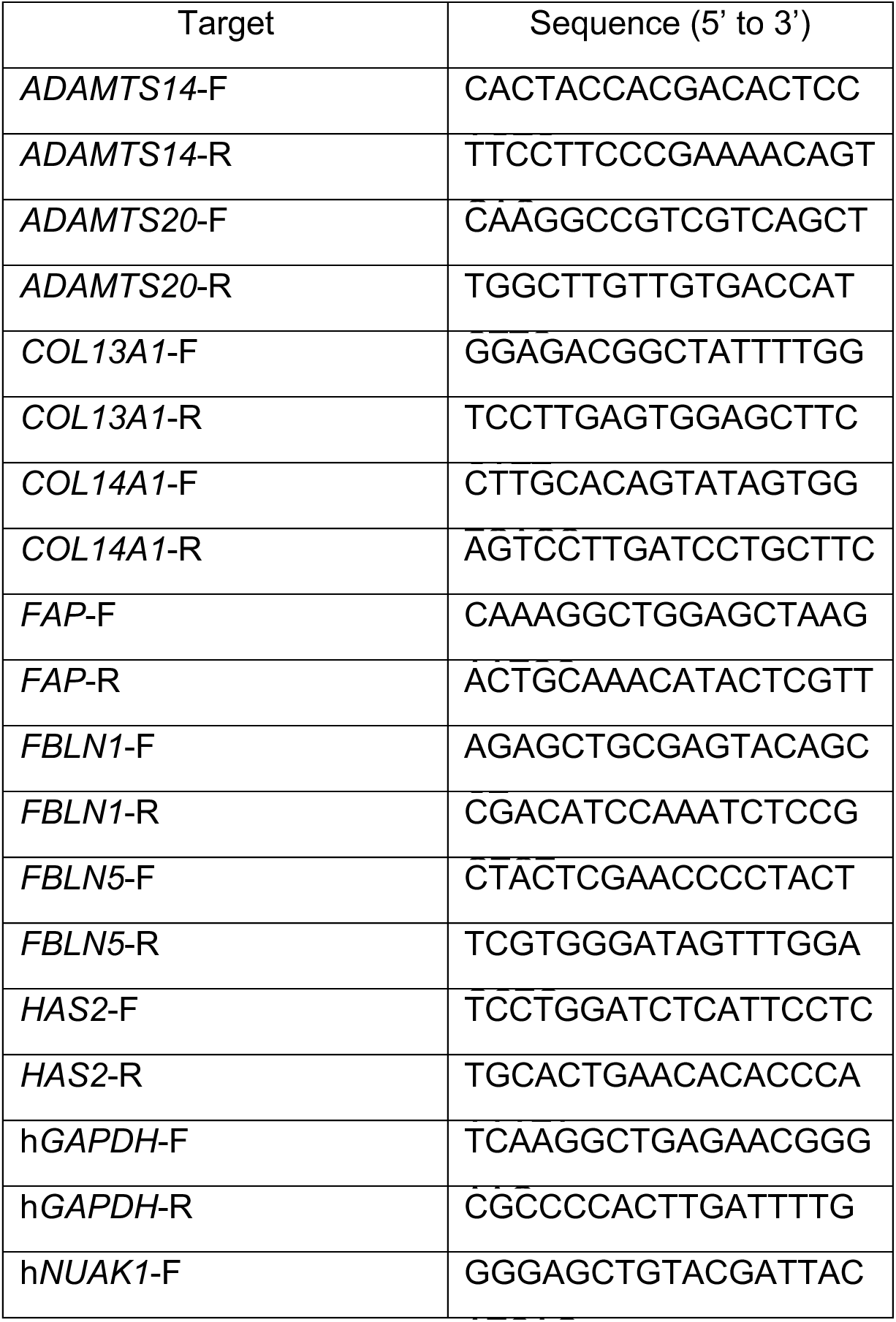
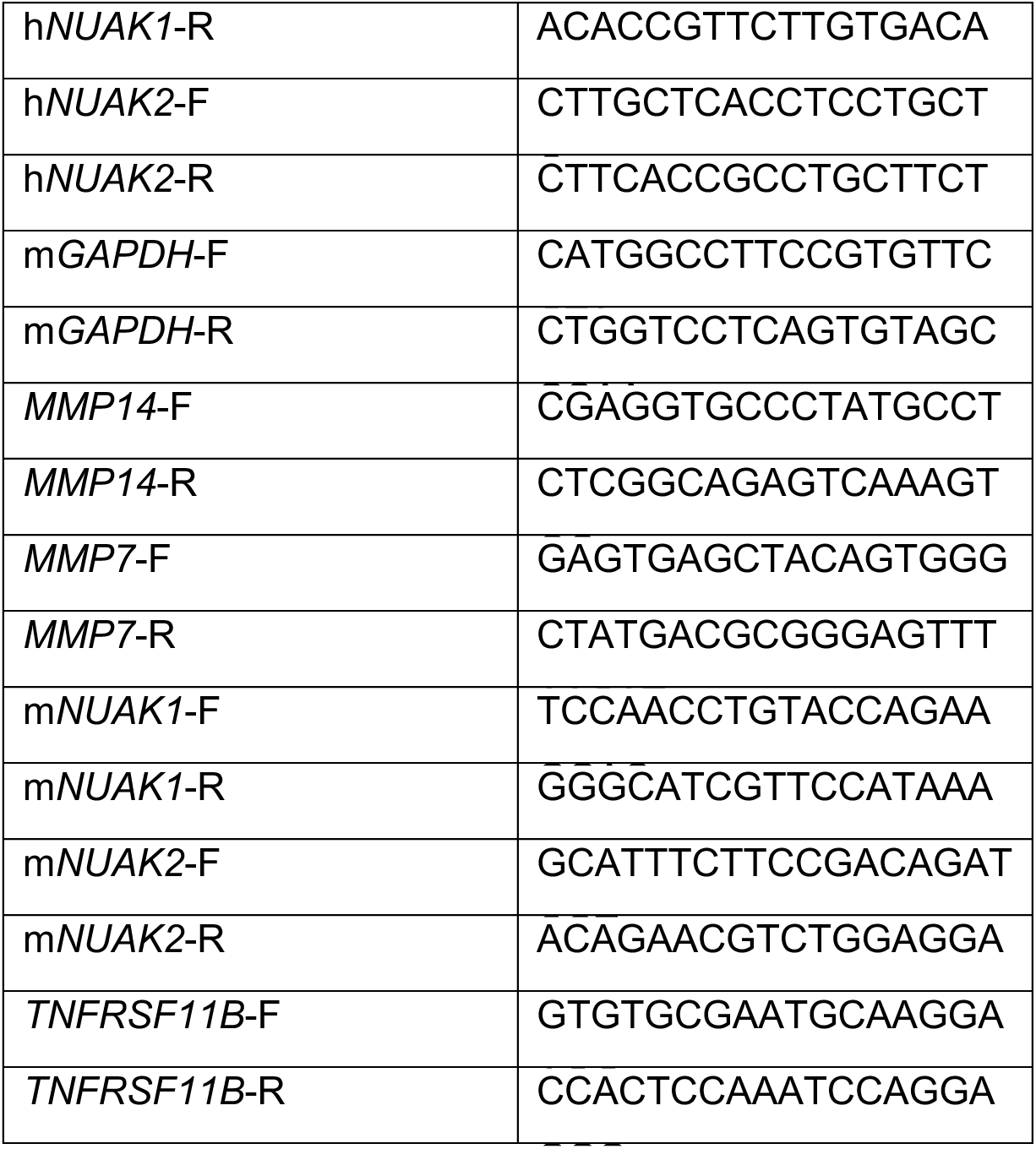
List of qPCR primers.

**Appendix Table S2.** List of Antibodies.

| **Name** | **Company** | **Cat. #** | **Application** |
| --- | --- | --- | --- |
| anti-Alpha tubulin | GeneTex | GTX628802 | WB (1:1000) |
| anti-E-Cadherin | CST | 3195 | WB (1:500) |
| anti-GAPDH | Millipore | MAB374 | WB (1:1000) |
| anti-Ki67 | CST | 12202S | IHC (1:500), ICC (1:500) |
| anti-N-Cadherin | CST | 13116 | WB (1:500) |
| anti-NUAK1 | CST | 4458S | WB (1:500) |
| anti-NUAK2 | abcam | ab224079 | WB (1:1000), ICC (1:100), IHC (1:50) |
| anti-Slug | CST | 9585 | WB (1:500) |
| anti-Snail | CST | 3879 | WB (1:500) |
| anti-Vimentin | CST | 5741 | WB (1:500) |
| anti-ZEB1 | CST | 3396 | WB (1:500) |
| anti-ZO-1 | CST | 8193 | WB (1:500) |
| anti-β-Catenin | CST | 8480 | WB (1:500) |

**Appendix Table S3.**
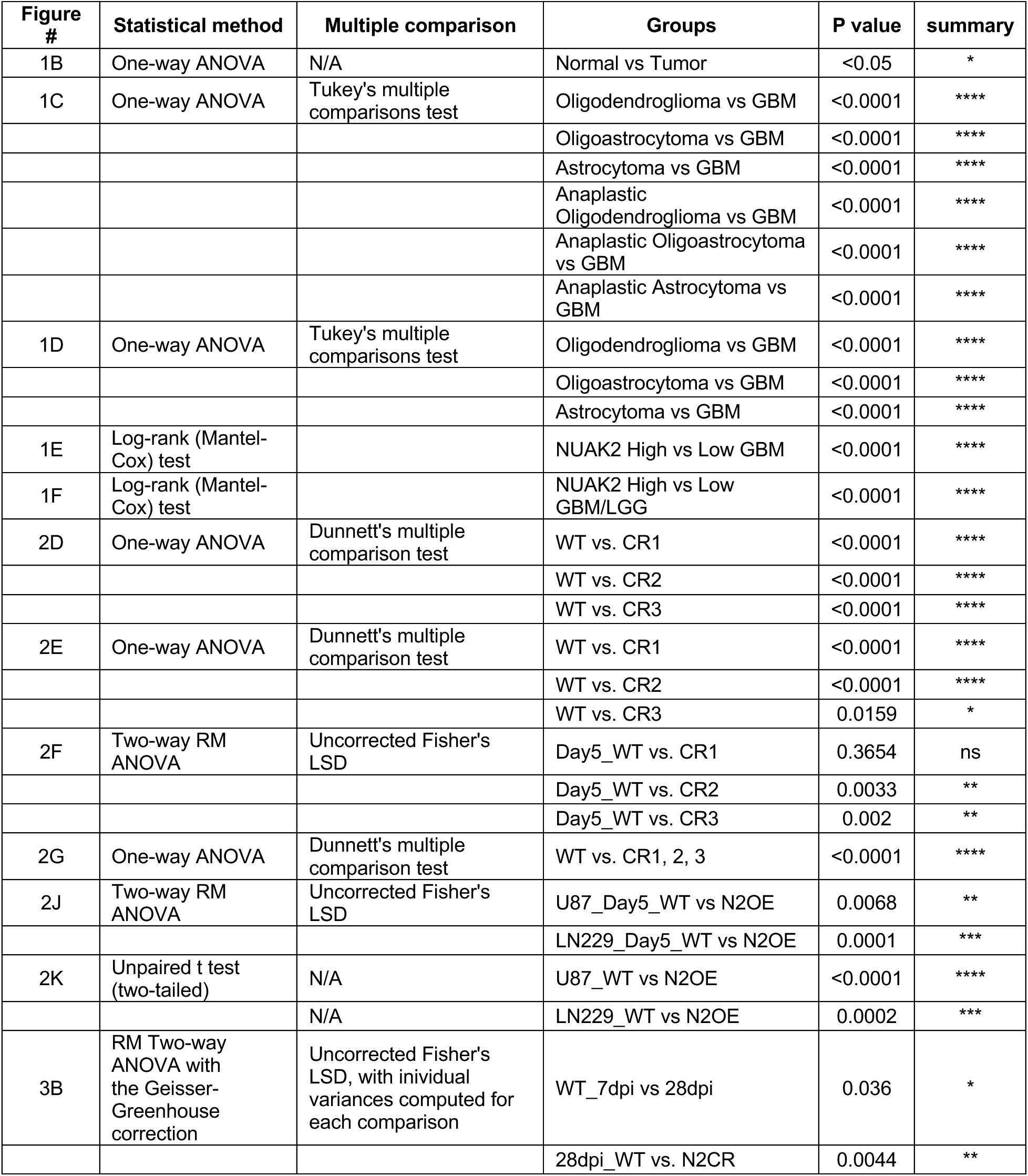
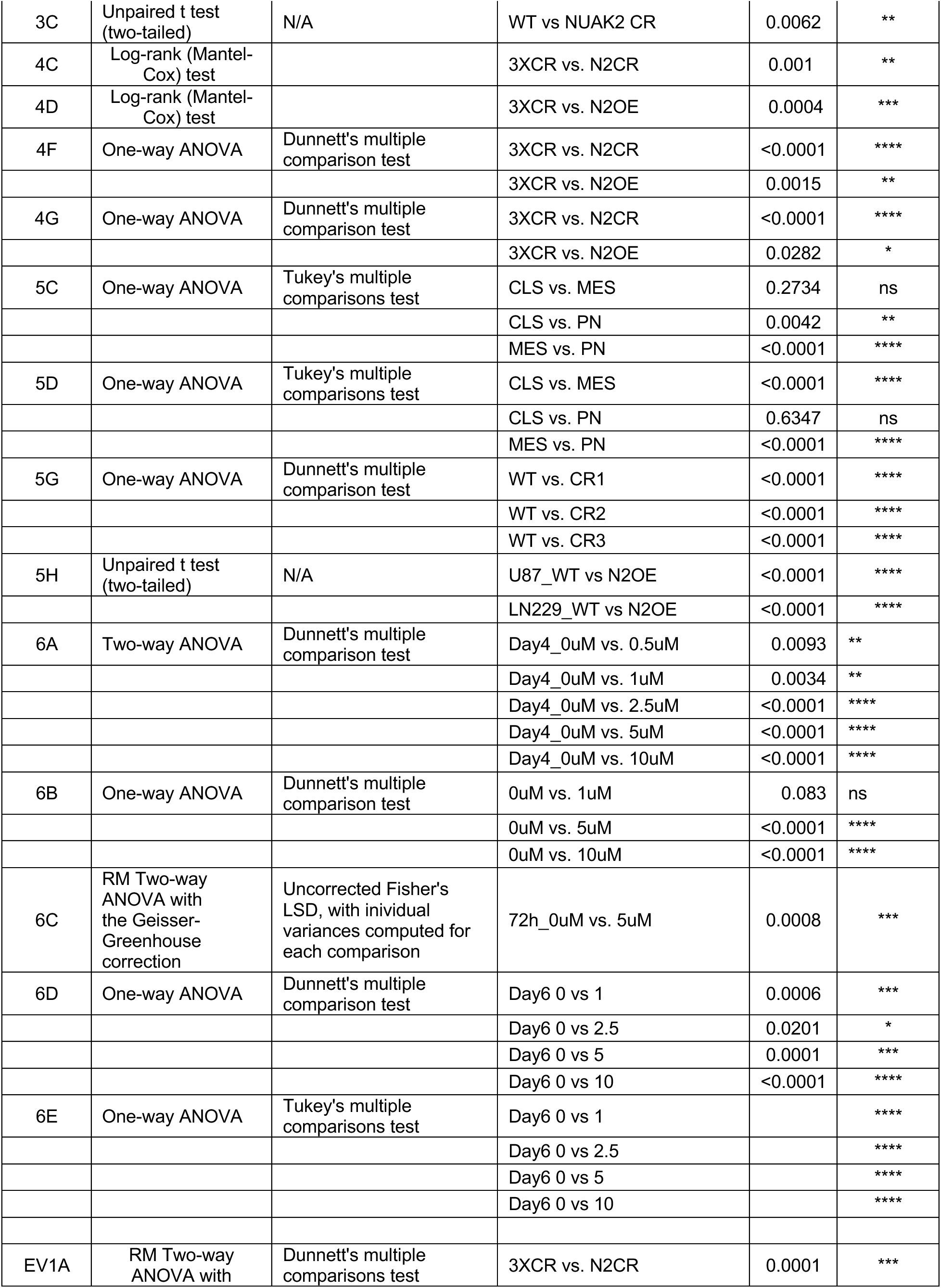
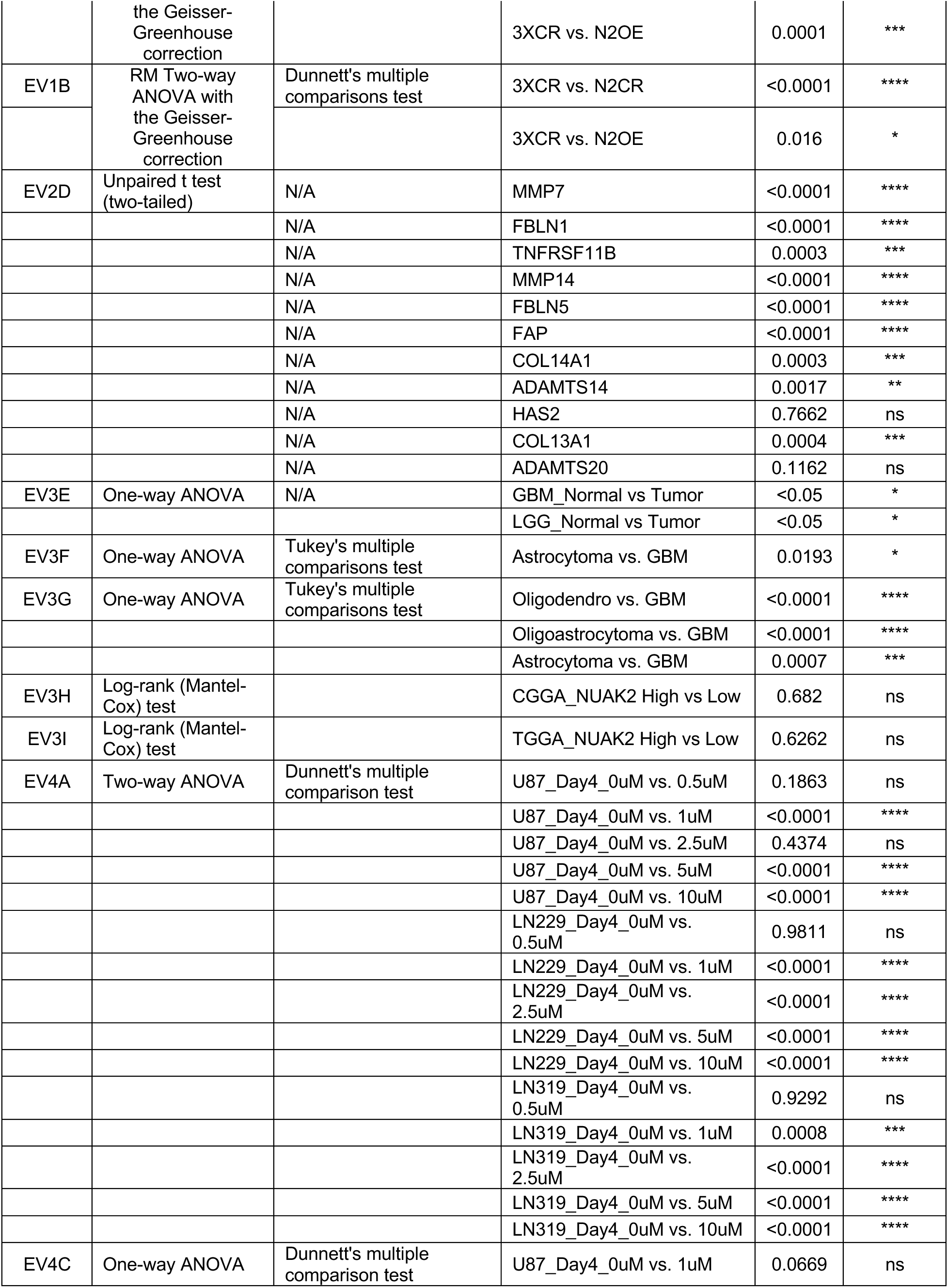
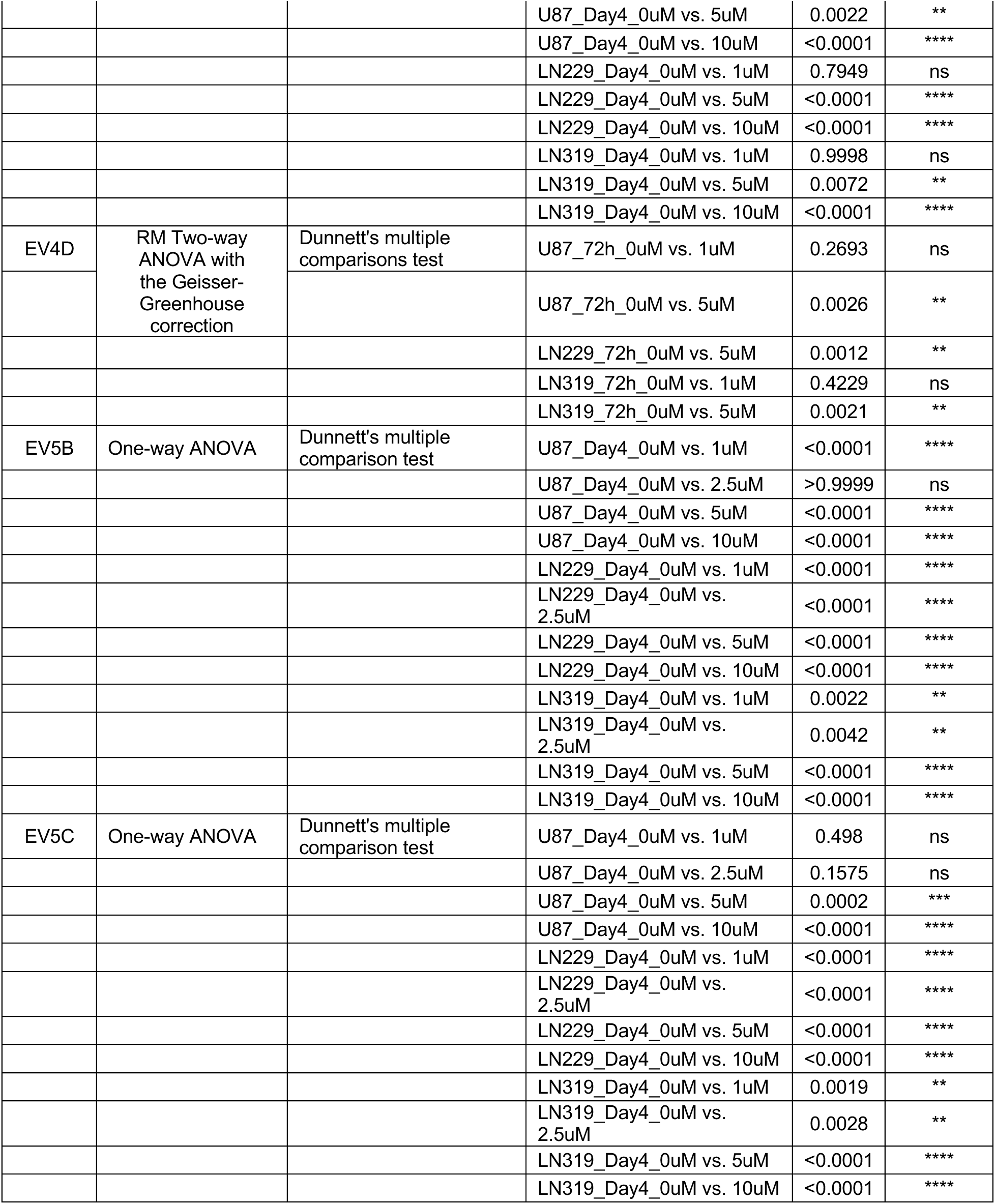
Summary of statistical tests and p values.

